# Olfactory performance explains duality of antennal architectural designs in Lepidoptera

**DOI:** 10.1101/2025.09.01.669131

**Authors:** Mourad Jaffar-Bandjee, Thomas Engels, Thomas Steinmann, Gijs Krijnen, Jérôme Casas

## Abstract

Male attraction by females through sex pheromones is widespread among Lepidoptera, and antennae are key olfactory organs during male orientation. Broadly speaking, two designs of antennae coexist in Lepidoptera: complex (pectinate) or stick-like (filiform) ones. Pectinate antennae have attracted attention because of their multiscale geometry, assumed to outperform filiform. Yet, the filiform design is by far more common. We compare the olfactory performance of the two designs using modeling, particle image velocimetry on 3D-printed scaled-up models, and computational simulations. In terms of absolute odor capture, pectinate antennae perform better at nearly all flying speeds. However, when considering drag, filiform designs are more energy-efficient than pectinate ones at low flight speeds, while the reverse holds at high speeds. This is due to the differential scaling of drag and molecule capture with flight speed. According to our results, small and slow moths would bear filiform antennae whereas big and fast moths would have pectinate ones, which is the general trend observed in Nature. We discuss exceptions to this general pattern and how species could evolve from one design to the other by investigating the influence of the antennal structural elements.

## Introduction

Many insect males detect chemical cues in the form of sex pheromone to find female mates, a step of paramount importance for species survival. The tiny amounts of sex pheromone emitted by females and the long-range orientation behavior of males make the moth communication system one of the most studied chemical communication system in the animal kingdom. [1, 7, 45]. Furthermore, sex pheromones are excellent tools in pest management for agroecology because they are highly specific to one insect species [40].

Once released in the air, pheromone travels either as single molecules or adsorbed on aerosols [32, 57] and must be collected by males, making chemical communication over long distances an interesting fluid-dynamics and mass transport problem. After emission from its source, atmospheric wind and turbulence inevitably stretch an odor plume into filaments and break it into isolated patches [51, 9]. At that level, searching for the odor source therefore requires sophisticated strategies such as zig-zagging and info-taxis [8, 35, 63]. Because antennae are the main organs of olfaction, one might postulate that the morphological design of antenna giving the best olfactory performance should be an evolutionary-selected attribute. Increasing the antennal surface area, and hence the number or size of sensors, seems *a priori* an obvious solution, a deduction already made by Darwin [14]. Surprisingly, the large group of Lepidoptera mainly exhibits two designs, the clavate and serrate forms being less extreme variants: filiform and pectinate antennae [43]. Other designs of complex architecture exist in other insect groups, from Hymenoptera to Coleoptera [46]. The relatively rare occurrence of the complex pectinate design in Lepidoptera is a second surprise. Recently, a phylogenetic study involving substantial fossil material concluded that antennae of complex architecture were not a key innovation, as they did not lead to the occupation of new ecological niches [49] nor to marked radiation [21].

Why the pectinate antennal design is not a key innovation, despite having evolved several times, might be due to the associated costs. A comprehensive research program, led by M. Elgar from the University of Melbourne, and a few other groups, has identified a large variety of costs associated with increasing length or structural complexity. They range from the costs associated with the neural information processing and maintenance of the many sensors, to the ability of the immune system to respond to challenging attacks or any investment in other body parts [69, 15, 56, 54]. The overall impression is that the cooccurrence of multiple designs might be the expression of a delicate balance between costs and benefits, precluding the cascading positive effects typical of key innovation. Such balance is well studied in vision for example [55, 26].

Experimentally comparing the two most extreme designs of filiform and pectinate antennae along those lines is hampered by the complex geometry of the pectinate antenna made of very delicate structures spanning many orders of magnitudes in size. So far, the fluid dynamics of pectinate antennae have been the object of many studies, either an experimental one [65] or through modeling [10, 28]. In the modeling works, the pectinate antenna geometry was strongly simplified. The study of only two hierarchical levels of a pectinate antenna with a more realistic geometry was achieved [30], but the embedding of these results into those of a complete antenna remained beyond reach, and with it an assessment of the enhanced olfactory performance of the whole organ.

The pectinate antenna is indeed a complex hierarchical structure with several levels [59]: the flagellum spanning the entire length of the antenna, the rami (secondary branches perpendicular to the flagellum) and the sensory sensilla located on the rami. There is a difference in size of almost four orders of magnitude between the diameter of the sensilla (3 µm) and the total length of the antenna (1 cm). Another difficulty is the high number of sensilla (tens of thousands), which are high aspect ratio cylindrical structures.

3D-printing micrometric cylinders with high aspect ratios is a key issue, as the materials used in printing technologies do not nearly reach the resilience of biological structures of similar sizes. Long and thin 3D-printed cylinders often result in very loose and fragile overall structures that may easily be deformed and damaged. Upscaling the structure could overcome this issue. However, the four orders of magnitude in antenna dimensions would result in a large structure that would be exceedingly long to print. For example, printing an antennal structure of several centimeters with details of 40 µm requires an estimated writing time of almost 3 days with one of the best tools (Nanoscribe Photonic Professional GT) and the results is very fragile due to the thin cylinders. In terms of numerical simulations, the high number of cylinder-like sensilla is the main problem because it requires a very thin mesh in a large volume, resulting in extremely computer-intensive simulations.

Our approach is to first derive a simplified model of pectinate antenna based on fluid dynamics consideration and validated by realistic computer-intensive simulations. Second, similarly to what has been done in [30], we combine numerical simulations and experiments on these simplified mockups to assess the olfactory performance of a pectinate antenna.

The aim of this work is to use such a combination to compare the efficiency of pheromone capture by architecturally complex pectinate antenna organs with the efficiency of the simpler and more common filiform antennae. To go beyond the different biological costs associated with long and complex antennae listed above, we quantify the aerodynamical drag of both structures and compare those costs and benefits. The quantity of pheromone captured per unit drag enables us to identify two conditions for which either design is preferred: flow speeds and pheromone relative diffusion speed, via the Schmidt number. It also enables us to rank the relative importance of the structural elements of pectinate antennae in the overall performance.

## Materials and methods

For odor sensing in general, independent of the actual antenna morphology, two parameters play an important role: the kinematic viscosity of air, *ν*_air_, and the pheromone diffusivity, *ν*_phero_. While the former is well known, the latter is associated with some uncertainty. We assembled a number of diffusivities for different moth sex pheromones (SI, Tables S1-3), yielding values from *ν*_phero_ = 3.61 *×*10^−6^ m^2^ s^−1^ to 7.01 *×*10^−6^ m^2^ s^−1^. To account for the uncertainty in determining *ν*_phero_ we extended its range to [1 *×*10^−6^, 10 *×* 10^−6^] m^2^ s^−1^. The uncertainty is due to the non-exhaustiveness of our list of pheromones and the approximated value of their diffusion coefficients based on a theoretical formula [64]. The ratio of air kinematic viscosity to mass diffusivity is called the Schmidt number, here Sc = *ν*_air_*/ν*_phero_, where Sc ∈ [1.6, 15.6] for the range of diffusivities considered here. Pheromone puffs are advected by the flow and follow its streamlines, but also diffuse in all directions and thus move perpendicular to the streamlines, which allows them to be captured by the antenna. Higher Sc thus implies that a pheromone is more difficult to capture.

Olfactory performance of both filiform and pectinate antennae was also investigated over a range of velocities. Moths usually fly between 0.5 and 3 m s^−1^ [2, 3, 30]. The range of velocities investigated here was extended to 0.1 to 5 m s^−1^. The antennae were considered to be perpendicular to the air flow. During flight, both types are most likely bent, but we are unaware of studies of the dynamical extent of bending at sufficient high spatial and temporal scale.

### Morphology of filiform and pectinate antennae

Our study is centered on the comparison of filiform antennae, which is the predominant antenna design in insects, with their pectinate counterparts (Fig. 1A-B). Filiform antennae occur with diameters 60–120 µm [67] and variable length, but to simplify comparison we use the same length as in the pectinate antenna.

**Figure 1.**
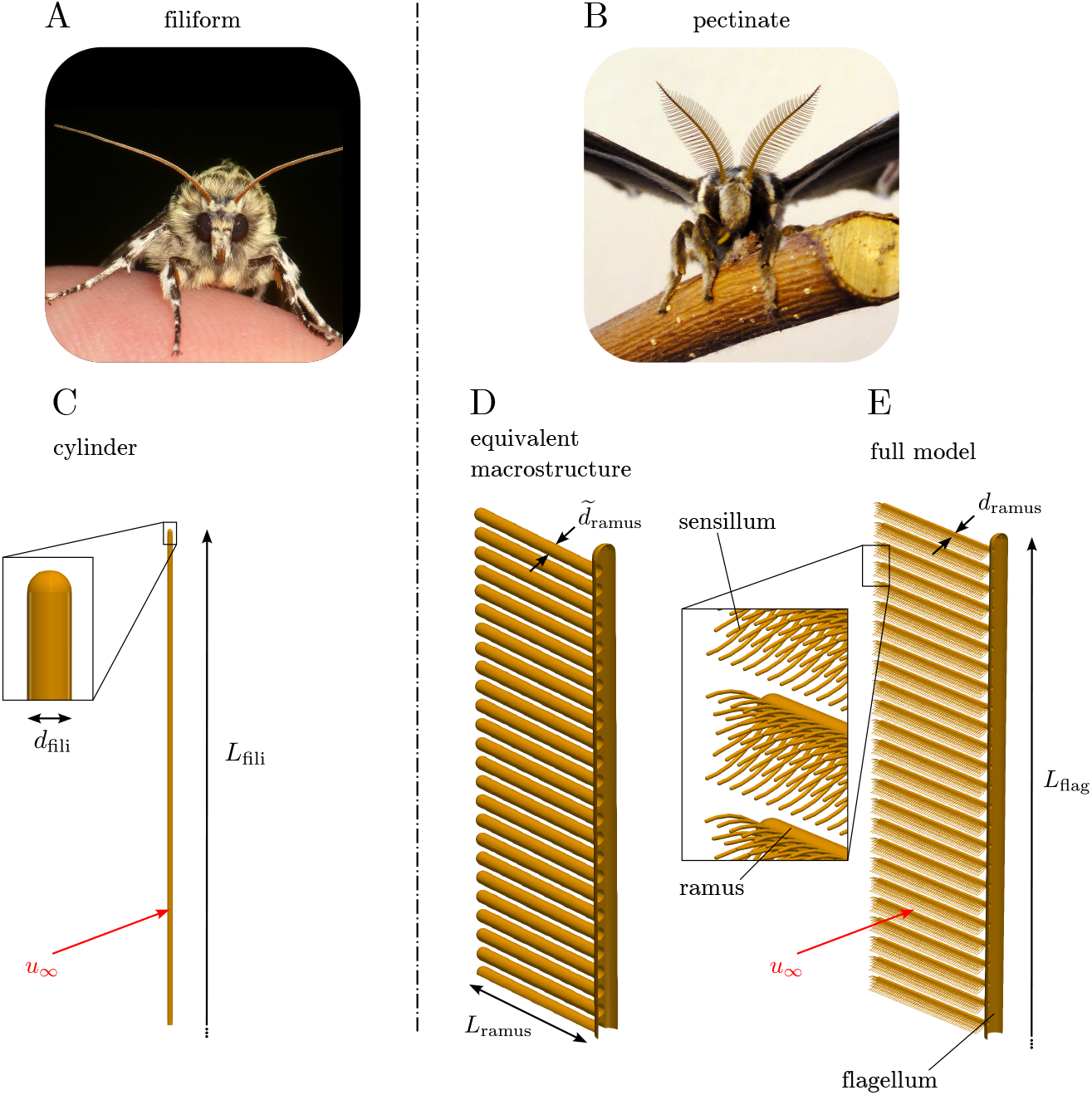
Morphology and modeling of filiform (A: Clearwing moth *Gaujonia renifera* (Noctuidae), photo by Andreas Kay) and pectinate (B: *Samia cynthia* (Saturniidae)) insect antenna. As the models are symmetric with respect to two planes, only a quarter is shown. See text for details.

**Table 1.**
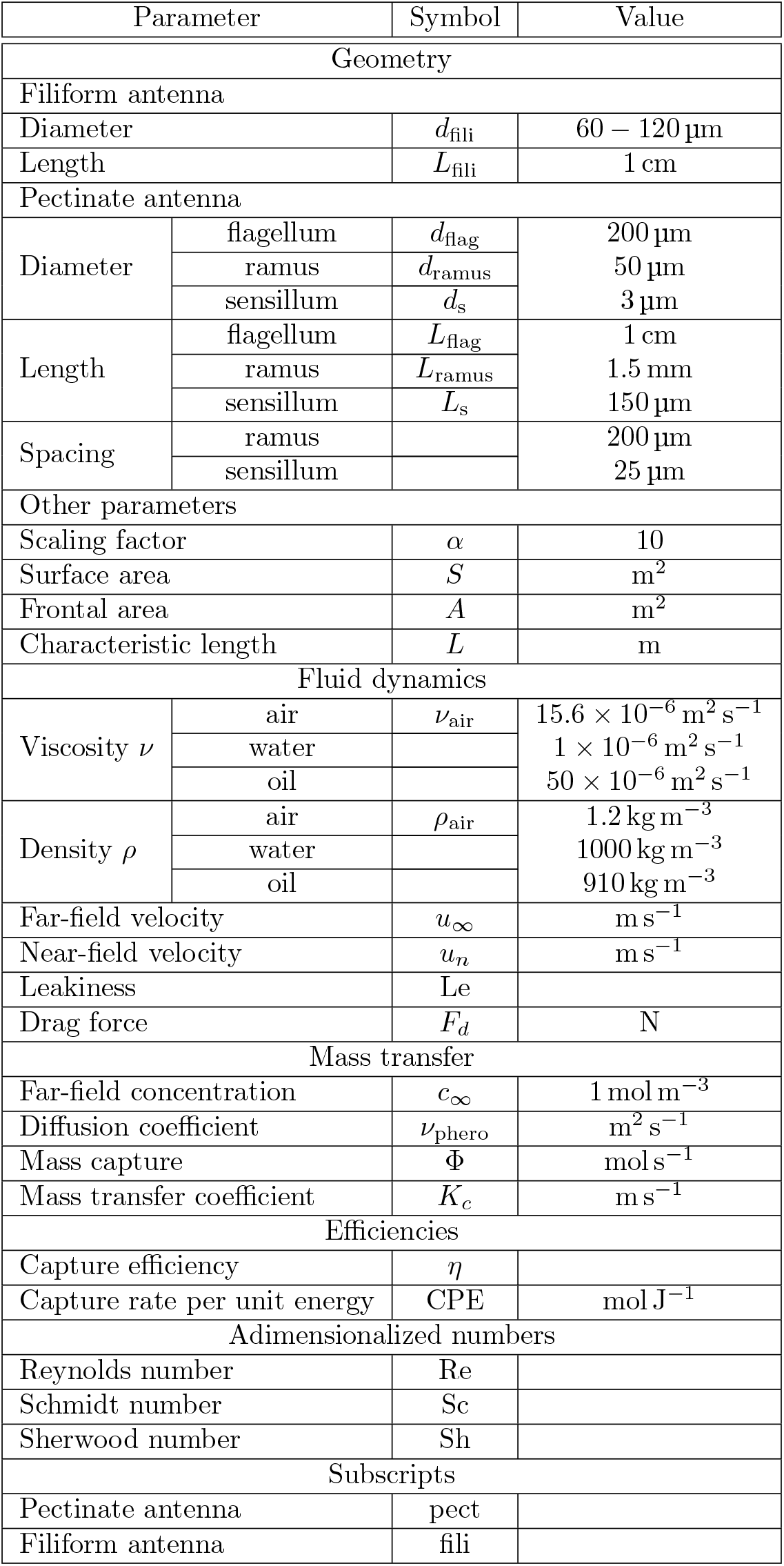
Parameters. The last two lin_6_es show the subscripts used to refer to respectively the pectinate antenna and the filiform antenna. For example, *η* is the symbol for the capture efficiency. *η*_pect_ refers to the capture efficiency of the pectinate antenna and *η*_fili_ refers to the capture efficiency of the filiform antenna.

Regarding pectinate antenna, we focus on one species, *Samia cynthia* (Saturniidae), because its antenna has already been studied [31, 30]. Pectinate antennae are multiscale objects consisting of three levels: a central stem (flagellomere), to which the rami are attached, each of which is finally covered with sensillae (Fig. 1E). The flagellomere is *L*_flag_ = 1 cm long with a diameter *d*_flag_ = 200 µm. It carries *N* = 100 rami, 50 on each side. For simplification we assume identical rami (*L*_ramus_ = 1.5 mm, *d*_ramus_ = 50 µm), spaced 200 µm apart, in our model (Fig. 1B). The sensillae, which are sensory hairs, are the tiniest elements (*L*_*s*_ = 150 µm, *d*_*s*_ = 3 µm). Each ramus carries *N* = 280 sensillae, arranged in groups of five with a spacing of 25 µm. Contrarily to what might be expected, sensillae are oriented towards the flow. We refer to a ramus and its sensillae as the <u>‘microstructure’</u> [30], and to the flagellomere and its rami (excluding sensillae) as the <u>‘macrostructure’</u>.

### Modeling of filiform antenna

We model the filiform antenna as a straight cylinder (Fig. 1C). As it is slender (*L*_fili_ ≫ *d*_fili_) we can neglect edge effects at tip and base; consequently, absolute values, such as total pheromone capture, vary linearly with *L*_fili_ and relative ones, such as efficiencies, are independent of *L*_fili_. We are interested in two aspects of the model: its aerodynamics and its pheromone capture. Because unsteady odor influx is a problem on its own that affects filiform and pectinate the like, our study is restricted to the steady state regime (see the next sections). The filiform antenna is also restricted to the steady state regime to facilitate comparisons.

<u>Aerodynamics</u> Defines the airflow around the cylinder and its drag, which is among the most often studied problems in fluid mechanics [68] (for more details, see SI).

<u>Pheromone capture Φ</u>, *i.e*. moles of pheromone captured per time unit, results from the interplay between airflow and pheromone diffusivity. It is defined as the flux of pheromone from the air to surface of the cylinder. The concentration at the surface of the cylinder is assumed to be zero. We extract Φ_fili_ from our 2D simulations and compare with analytical results from the literature. Throughout this work, we set the far-field concentration *c*_∞_ to 1 mol m^−3^. Real pheromone concentrations are much lower. However, the governing advection-diffusion equation is linear so the choice of an arbitrary far-field concentration does not alter our understanding of the mechanisms of pheromone capture. Analytically, we calculate it as

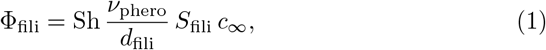

where *S*_fili_ is the cylinder’s surface. The Sherwood number Sh = *K*_*c*_*d*_fili_*/ν*_phero_ is the ratio of convective to diffusive mass transport and analog to the Nusselt number in heat transfer. It can be empirically calculated [42] from Reynolds-and Schmidt number as

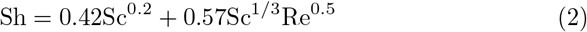

### Modeling of pectinate antenna

In order to cope with the multiscale nature of the pectinate antenna, we derive three distinct models from the original geometry: the equivalent macrostructure (Fig. 1D), the full model (Fig. 1E) and the pheromone capture model (Fig. S2). All models are restricted to the steady-state regime because unsteady effects in odor influx are a distinct problem and affect both designs. We also use the filiform model to correct the pheromone capture of the pectinate antenna at low velocities.

Pectinate antennae have usually leaf-like shapes. Here, the outline of the antennae was chosen rectangular. The main reason for this simplification is that it has been shown that most of the rami have a similar length [31], except at the tip. Moreover, this simplification decreases the number of parameters to investigate and allows to focus on the effect of ramus and sensillum lengths on the olfactory performance. Secondary reasons are that, with a rectangular shape, 2D velocity fields could be measured experimentally (see section ‘Experiments’) and, in the simulations, the model could be reduced to one quarter thanks to symmetry boundary conditions (see sections ‘Simulations’ and ‘Full model of pectinate antenna’). The reduction of the model size was especially important for the full-model simulations (see section ‘Full model of pectinate antenna’) which already required two millions CPU hours (see SI).

### Equivalent macrostructure model

The pectinate antenna as a whole spans four orders of magnitude in size. While numerical simulations including all sensillae are indeed possible, they are very expensive and cannot be used for a large parameter sweep of *u*_∞_ as is required for this study; they can be used only as validation. We therefore developed an equivalent macrostructure (Fig. 1D) where the effect of sensilla is taken into account by increasing the ramus diameter until it produces, without sensilla, the same drag as the original rami with sensilla (for more details, see SI). Such an approach was already used on permeable dandelion seeds [13], which were modeled by impermeable cylinders with equal drag, as well as in a study on bristled wings of beetles where hair-like structures with outgrowths were modeled by cylinders with appropriate diameters [41].

As in the case of the filiform antenna, we are interested in the <u>aerodynamics</u> and <u>pheromone capture</u> of the pectinate antenna. However, the equivalent macrostructure serves as a proxy for aerodynamics only. In addition to the drag force, which is also present for filiform antennae, pectinate antennae share a common feature with all porous structures, the leakiness, which measures how much air goes through the antenna. We define it as the portion of the airflow going through the antenna compared to the far-field air flow [30],

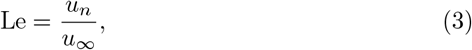

where *u*_n_ is the mean velocity inside the antenna. Leakiness generally increases with increasing *u*_∞_; at low velocities however, rake-like structures such as our macrostructure model can behave aerodynamically almost as an impermeable solid plate [19, 11, 17].

We determine drag and leakiness for the equivalent macrostructure model using simulations and experiments in a steady-state regime.

#### Simulations

We run 3D simulations (Comsol v5.6) to determine the fluid dynamics around the *<u>equivalent macrostructure</u>*. We restrict the simulations to a quarter of the model and use appropriate symmetry boundary conditions to reconstruct the entire structure (for more details, see Fig. S5). We investigate a range of velocities and determine the steady-flow solution. To obtain Le, we integrate the flux of air between the rami and determine the average velocity *u*_*n*_ through the equivalent macrostructure at each velocity *u*_∞_. The simulations also provide the drag force. More details of the simulations are available in the SI.

#### Experiments

For PIV experiments, we use the same setup and method as in [31]: we measure velocity fields around scaled-up equivalent macrostructures in oil or water using appropriate velocities to ensure that the Reynolds numbers are kept constant. We then obtain the drag force *F*_d_ from the PIV measurements through a balance of momentum, assuming the flow field is 2D (for more details, see SI).

### Full model of pectinate antenna

Our full model includes 100 rami and 28000 sensilla (Fig. 1E), and we study it only in numerical simulations. Commercial codes can currently not handle such multiscale geometries, and we resort to the open-source code WABBIT [16] for such simulations. Its key feature is automatic refinement of the numerical grid using wavelets, combined with an efficient parallelization on large-scale supercomputers. Wavelets allow refining the grid where it is necessary to ensure a given precision, while in other regions coarser resolution is used. We exploit the two symmetries and simulate only a quarter of the full model. Simulations are run until the steady state develops. To give an idea of the complexity, we used 4096 CPU cores and 8 TB of memory and a simulation typically involves 4 billion grid points. Even with this sophisticated approach, we could not simulate the animals’ geometry unless we increased *d*_*s*_ to 6 µm – we verified that the impact on force and leakiness is negligible. Still, because of the tremendous cost, we simulate the full model only for three values of *u*_∞_, yielding Le and *F*_d_. The full model simulations thus serve as validation for the assumptions made to derive the equivalent macrostructure. More details on the full model simulations can be found in the SI.

### Pheromone capture model of pectinate antenna

Using the equivalent macrostructure and the full model, we determined leakiness Le and drag *F*_d_. We now turn to the actual capture of pheromone. Similarly to the filiform antenna, the capture of pheromone is defined as the flux of pheromone from the air to the surface of the antenna per unit of time. Pheromone absorption takes place mainly in the sensilla [38]. To compute the capture rate of the pectinate antenna, Φ_pect_, we consider that at close range, the network of rami and sensilla can be considered as an infinite array of infinitely long rami, all bearing sensilla, in the exact same way as what we do to calculate the diameter of the rami in the equivalent macrostructure. We then run steady-state mass transport simulations (Comsol v5.6) on a section of ramus with five sensilla and with appropriate periodic conditions to mimic an infinite array of infinitely long rami. The section of ramus is 25 µm high and the inter-ramus space is 200 µm. The far-field pheromone concentration is set to 1 mol m^−3^ and the concentration at the surface of the sensilla is considered zero. This model does not consider diffusion of pheromone from the air flowing around the antenna. At low velocities, this may be a problem so we applied a correction on the amount of captured pheromone (for more details, see SI.)

### Capture Efficiency and Capture-Per-Energy

We define capture efficiency *η* as the capture rate normalized with the free stream pheromone flux through the projected area *A*,

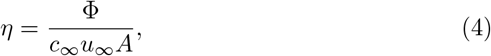

where A is defined by *A*_fili_ = *d*_fili_*L*_fili_ for a filiform antenna and *A*_pect_ = 2*L*_flag_*L*_ramus_ for a pectinate antenna *A*_fili_ = *d*_fili_*L*_fili_ and *A*_pect_ = 2*L*_flag_*L*_rami_. By construction, *η* removes the dependency on the surface *A*, which makes both antenna designs more comparable. However, from the point of view of an animal, the absolute capture rate Φ is what matters. Note *η* can exceed 100% if pheromone is entrained from the sides (cf above). As the normalization with a reference flux is somewhat arbitrary, we use a second measure to compare both antenna designs: the capture rate per unit energy (CPE). It is defined as

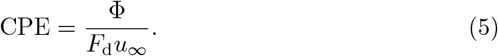

It thus puts the capture rate in relation to the power that has to be spent to carry the antenna, which is the product of its drag and velocity. This measure is similar to the lift-to-power ratio [18] sometimes used to characterize flight energetics.

## Results

### Leakiness

We begin with the leakiness of the full model of the pectinate antenna. Fig. 2A-B shows the normalized axial velocity passing through the antenna for the 1 and 5 m/s far-field velocity cases. The effect of reduced velocity is clearly visible: the same structure behaves more like a solid plate in the low-velocity case, with little fluid passing through it (10%). By contrast, at higher inflow velocity, Le is significantly higher with the inter-ramus flow velocity reaching almost *u*_∞_: in total, almost 40% of the air pass through the structure (Fig. 2C). For pheromone sensing, it means that at high velocity, a large pheromone flux is available, but it also has less time to diffuse to the antenna surface. It is these two competing processes that determine the performance of the pectinate antenna [30].

**Figure 2.**
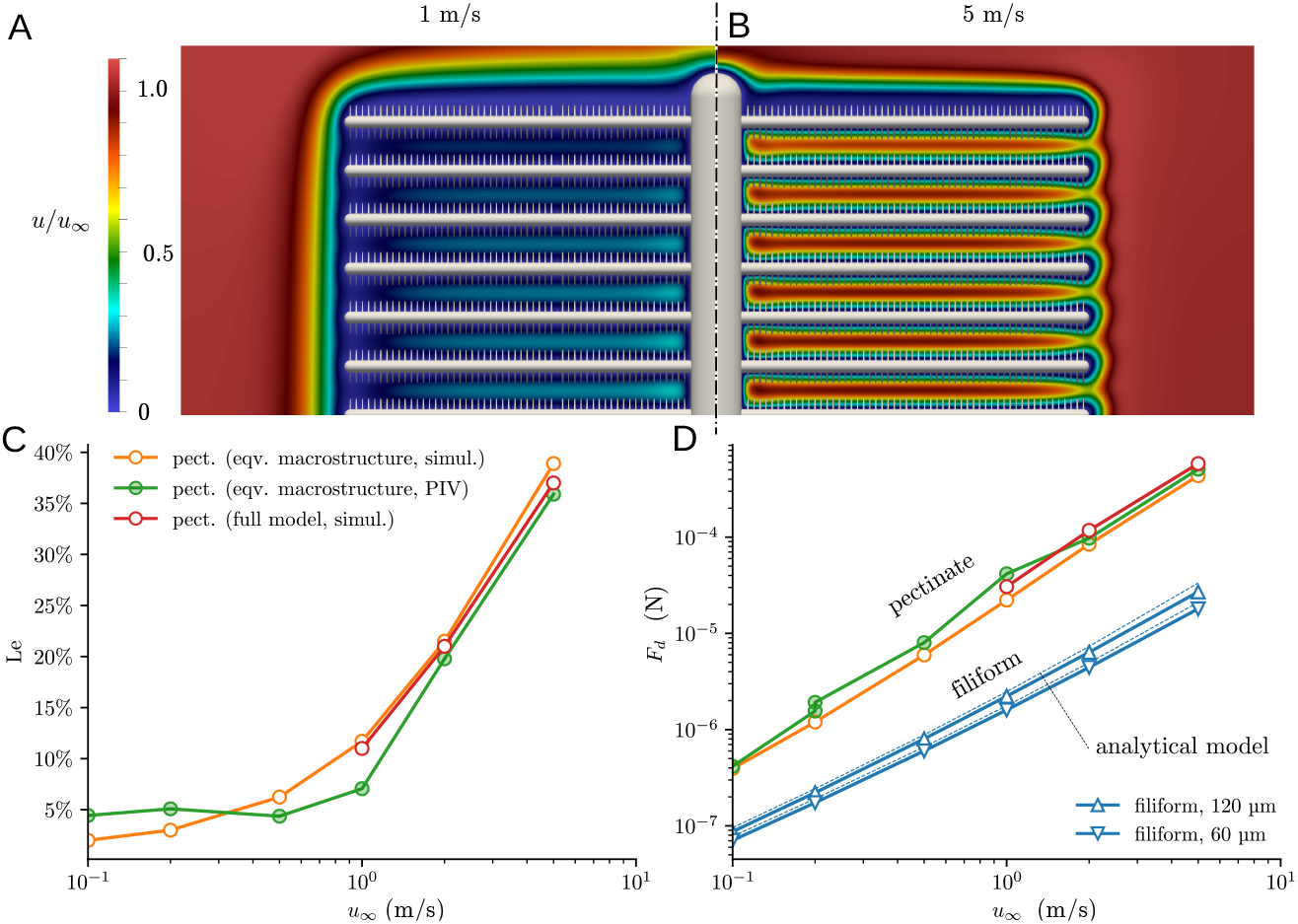
Leakiness of full model and equivalent macrostructure. A-B: axial velocity (*u/u*_∞_) of the fluid passing through the full model antenna, where *u*_∞_ = 1 m/s (A) and 5 m/s (B). These data are obtained numerically using WABBIT. C: Comparison of leakiness of equivalent macrostructure with the leakiness of the full model. Equivalent macrostructure data from numerical simulation (COMSOL) and experiments (PIV). Both curves agree well with the full model leakiness obtained from numerical simulation (WABBIT). D: drag force *F*_d_ for the pectinate models and the filiform antenna.

Figure 2C compares the leakiness obtained by our three methods. The full model simulations yield the most faithful results but are limited to three velocities because of the numerical cost. The numerical simulations of the equivalent macrostructure, where sensilla are modeled by an increased ramus diameter, yield very similar leakiness. This proves that the concept of the equivalent macrostructure is a suitable way to reduce the geometrical complexity of the problem, as it yields by construction the same drag and very similar leakiness. The numerical data are furthermore supported by our PIV measurements, performed on the equivalent macrostructure. Because of the good agreement, we use the leakiness determined from the equivalent macrostructure simulations in the following.

In order to test the validity of our approach on a different structure, we apply the method to a grid composed of two rows of cylinders perpendicular to each other. Using Comsol, we determine both the leakiness of the grid and the leakiness of the structure with equivalent cylinders and found that they are indeed close to one other (for more details, see the SI).

### Drag force

Pectinate antennae are subject to higher drag than filiform ones, with a difference around one order of magnitude (Fig. 2D). In order to assess the biological importance of the cost of the drag in the moths energy budget, we compare the power lost in the drag of the two antennae and with the total energy consumption of the moth during flight.

Metabolic rates of insects during flight have been measured several times on moths (Sphingidae and Saturniidae [4, 25], *Spodoptera* [53]) and other insects (bees [61]) and are of the same order of magnitude, *V*_*O*2_ between 50 and 150 cm^3^*O*_2_/g/h (oxygen consumption per gram and hour). The average mass *M* of a male of *Samia cynthia* is around 1.5 g [39], and the relationship between mass and oxygen consumption is *V*_02_ = 59.43 *M* ^0.818^ [4]; this gives an average oxygen consumption of 83 cm^3^*O*_2_/g/h for a male of *Samia cynthia* in flight.

Oxygen consumption can be expressed in terms of metabolic energy consumption using the conversion factor 20 J/cm^3^*O*_2_ [5], leading to a total energy consumption of 0.46 W. Mechanical efficiency of flying moth is also around 10% [60], so the power spent on flying is reduced to 0.046 W. We divide the power lost in the antenna drag by the metabolic power spent on flying to assess the importance of the drag of the two antennae of a moth. At high velocity (5 m s^−1^), the power spent in the pectinate antennae is not negligible, reaching almost 10% of the total metabolic power consumed by the moth. In the case of a 10 mm long filiform antennae, large diameter antennae generate a higher drag compared to thinner ones but the relative power spent in the drag is still very low, reaching only 0.6% at most.

The metabolic costs of bearing pectinate antennae is comparable to the metabolic costs of other sexually-selected traits having an influence on locomotion. For example, swordfishes’ long swords increase the metabolic costs of swimming by around 20% [6] and hummingbirds’ tail the metabolic costs of flying up to 11% depending on the flight speed [12]. In these cases, the additional cost is due to the drag. But the additional metabolic cost can also be the consequence of an increased mass: in the case of stag beetles, the mass of their enlarged mandibles and their associated muscles increases the metabolic cost of flying by 26% [22]. In our case, the metabolic cost of the pectinate antenna is most likely underestimated because only the drag due to the antennae was considered here and not the maintenance costs of the neural cells of the antennae. These later costs are generally important in sensory organs [55]. In the case of the visual system of the fly *Lucilia cuprina*, they represent 10% of the metabolic rate at rest [27].

### Pheromone capture

The olfactory performance of filiform and pectinate antennae is shown in Fig. 3. We consider first the absolute capture Φ (Fig. 3A). For a given Schmidt number, the absolute capture rate of the pectinate antenna is always higher than the filiform one, regardless of the inflow velocity. In general, Φ increases with increasing *u*_∞_. Φ_fili_ is well approximated by a power law, yielding almost a straight line in the log-log diagram. As expected, increasing the Schmidt number makes it more difficult to capture the pheromone; increasing *Sc* by a factor ten results in a reduction of Φ_fili_ by approximately the same factor. We observe excellent agreement between our numerical simulations and Φ_fili_ from the analytical model (eq. 1). The pectinate antenna follows a largely different trend. For intermediate *u*_∞_, Φ_pect_ is virtually independent of the Schmidt number, while for low and high *u*_∞_, a reduction in capture rate is observed with increasing Sc.

**Figure 3.**
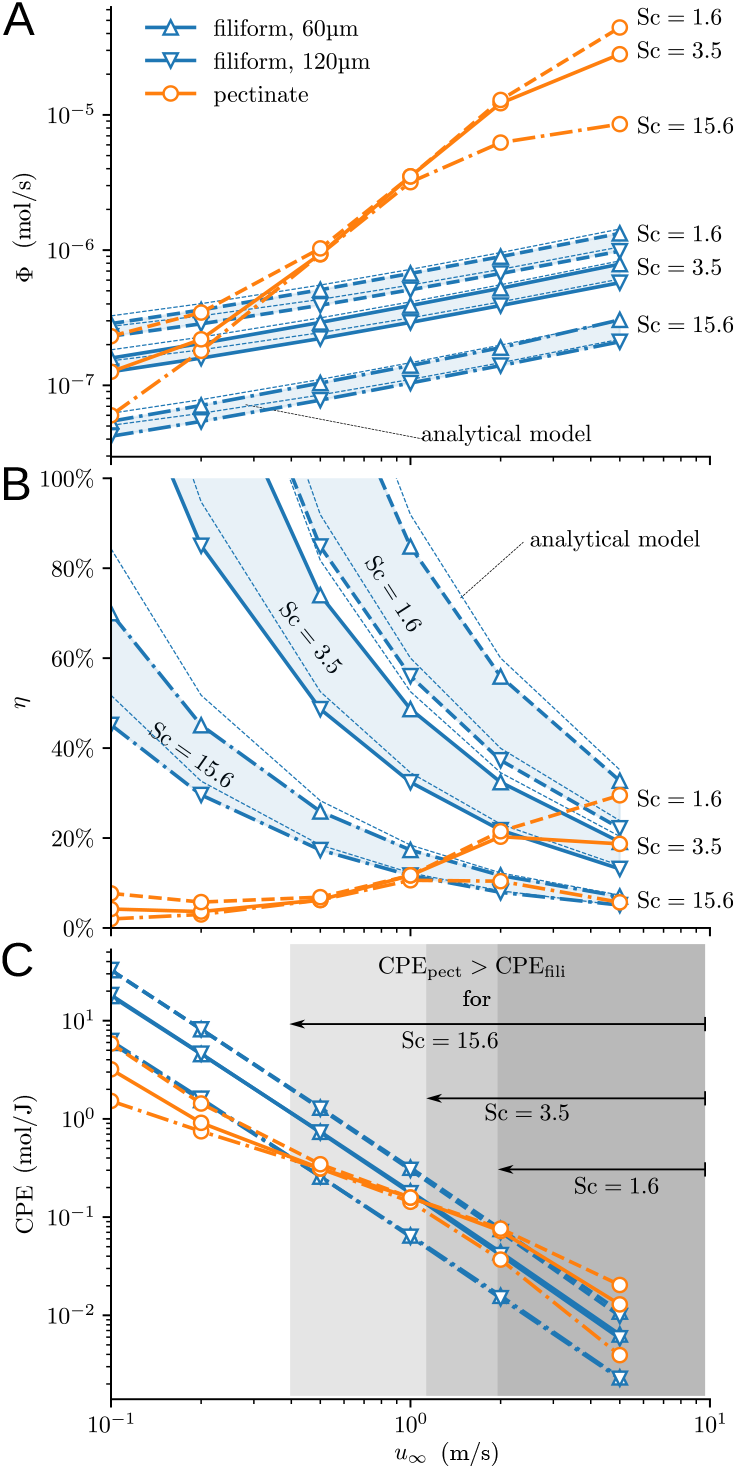
Pheromone capture of filiform (blue colors) vs. pectinate antennae (orange) as function of *u*_∞_. A: absolute capture rate (in mol s^−1^) of both antenna models. In absolute numbers, pectinate antennae capture much more pheromone molecules. Both models are computed with three different Schmidt numbers covering the range of various sex pheromones (represented by different line styles). For the filiform antenna, cylinder diameter ranges from *d*_fili_ = 60 µm to 120 µm (blue shaded surface) B: capture efficiency *η* (eq. 4) for filiform and pectinate antennae. Data for pectinate antenna are obtained by combining equivalent macrostructure leakiness with the pheromone capture model. C: capture rate per unit energy (CPE). Gray shaded areas mark the crossover of CPE for the different Schmidt numbers: for velocities in the gray area, the pectinate antenna outperforms filiform design.

The capture efficiency *η* (Fig. 3B) yields a different picture. Normalized by the reference pheromone flux, the filiform antenna is more efficient, except for the highest velocities tested. For the smaller Schmidt numbers, *η*_fili_ even exceeds 100% at low velocities: here, the pheromone entrainment from the sides becomes significant. By contrast, *η*_pect_ is rather low. This implies that for identical projected surface (cross-section), the filiform antenna is more efficient than the pectinate one - however, to obtain the same surface, *L*_fili_ would scale in meters rather than centimeters.

The capture-per-energy (CPE), shown in Fig. 3C, is a more suitable measure to compare both designs. In both cases, the energetic cost of carrying the antenna (*u*_∞_*F*_d_) increases faster than the capture rate Φ - consequently, CPE is monotonically reduced with increasing velocity. In the filiform case, CPE follows again a power-law behavior, but the CPE for the pectinate antenna behaves differently. For a fixed Schmidt number, at high velocities, CPE_pect_ is larger than CPE_fili_ by a factor 2, while at low velocities, the reverse holds true. This fact implies that for any given Schmidt number, there is a critical velocity above which the pectinate antenna outperforms the filiform one. This range is illustrated by the gray areas in (Fig. 3C). It depends on the Schmidt number: the higher it is, the more advantageous is the pectinate design. In other words, pectinate antennae are particularly efficient for pheromone that have a low diffusion coefficient, meaning they are difficult to capture, as well as at elevated velocities.

### Design parameters of pectinate antenna

We run additional simulations on Comsol to estimate the relative importance of two design parameters on the olfactory performance: the lengths of rami and sensilla.

We calculate the influence of ramus length on the leakiness, the drag, the pheromone capture and the capture-per-energy (Fig. 4, left column). The ramus diameter is 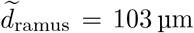 and we vary *L*_ramus_ as well as *u*_∞_. Leakiness is sensitive to ramus length. As *L*_rami_ increases, the leakiness increases as well. This phenomenon is most likely due to the relative importance of the flagellum, the thickness of the boundary layers and the edge effects. At high velocities, the boundary layers around the rami are reduced and interact less with each other. The flagellum, thicker than the rami, does have a boundary layer and is not permeable. Its relative importance compared to the surface of the antenna increases with the decrease of the ramus length. However, the main effects of an increase in ramus length are a linear increase in both the drag and the pheromone capture. As a result, the CPE varies little with ramus length. This is graphically for CPE.

**Figure 4.**
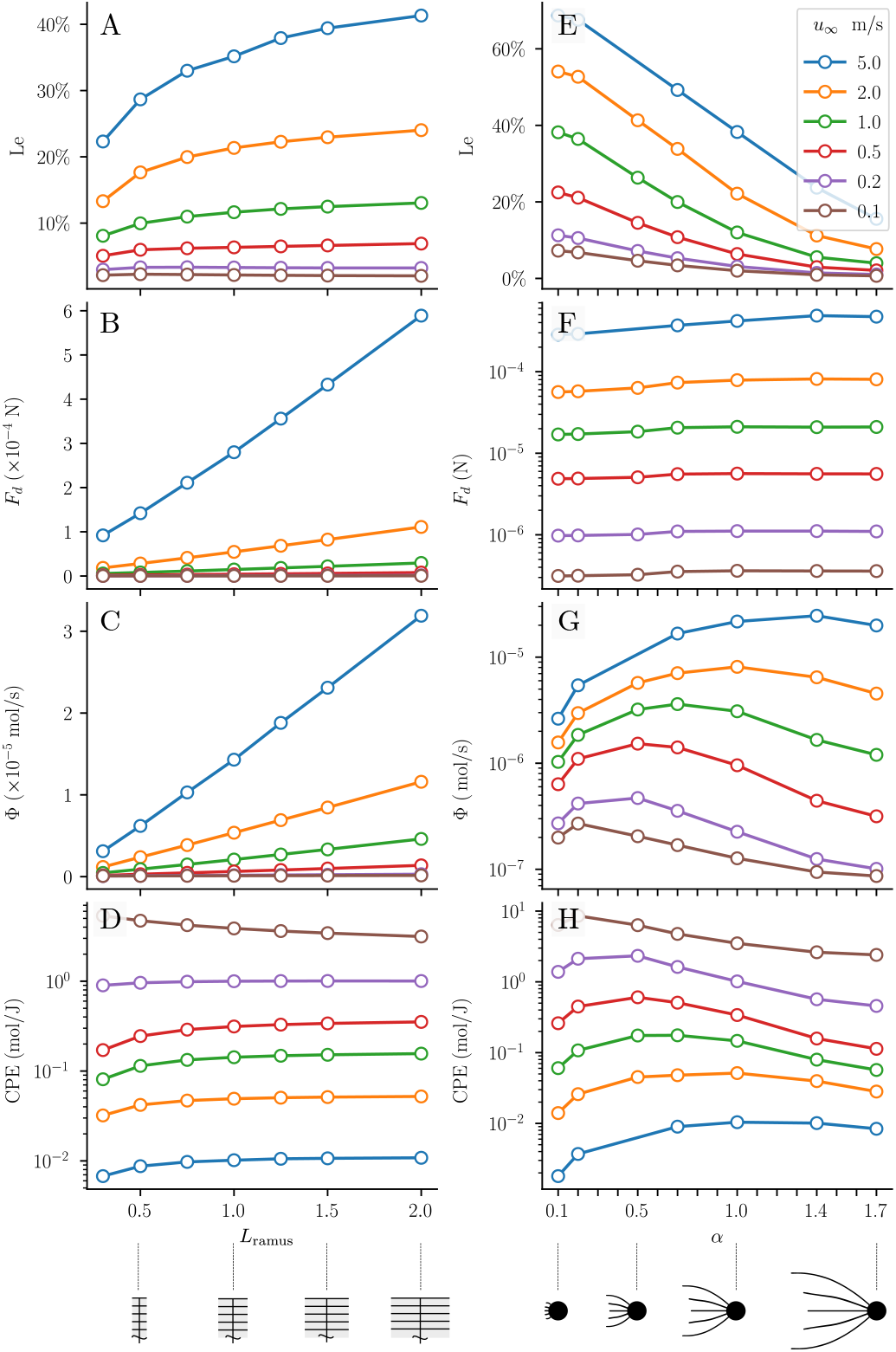
Influence of ramus and sensillum lengths on the olfactory performance of a pectinate antenna. Left column: influence of ramus length on (A) leakiness, (B) drag force, (C) capture rate and (D) capture-per-energy. Right column: influence of sensillum length on (E) leakiness, (F) drag force, (G) capture rate and (H) capture-per-energy. *α* is the scaling factor used to change the length of the sensilla but their diameter remained equal to 3 µm. *α* = 1 means the sensilla have the actual size of the moth sensilla. *α* 1 means the sensilla were scaled down and *α >* 1 means the sensilla were scaled up. Each color represents a free-stream velocity.

To determine the influence of the sensillum length, we run similar simulations to the ones done to determine the diameter of the rami in the equivalent macrostructure but, this time, we vary the length of the sensilla by scaling them with a factor *α. d*_ramus_ = 50 µm and *d*_*s*_ = 3 µm are kept unchanged. We find that the drag per unit length of such a structure changes with the length of the sensilla and calculate accordingly the diameter of the pseudo-rami used to model the microstructures. The sensillum length has a strong influence on the leakiness but, surprisingly, little effect on the drag (Fig. 4, right column). The effect on the pheromone capture is more complex. The pheromone capture increases first with sensillum length because it gives more time to the pheromone molecules to diffuse to the surface of the sensilla. Second, as the sensillum length becomes too long, the leakiness decreases and not much pheromone is available to the sensilla, implying that the pheromone capture decreases. The transition between the two trends, which also corresponds to the maximum of capture, depends on the far-field velocity. At low velocities (Fig. 4, right column, third line, violet and brown curves), the maximum of capture is reached for short sensillum lengths. At high velocities (Fig. 4, right column, third line, blue and orange curves), the location of the maximum is shifted to higher values of sensillum length.

Ramus length and sensillum length have thus different effects on the performance of a pectinate antenna. Increasing ramus length increases almost linearly the pheromone capture. However, the cost is that the drag also increases with ramus length almost linearly. Sensillum length has a less dramatic effect, for example on the drag, but it can help to fine-tune the performance of an antenna, especially at high velocities.

## Discussion

Our model of pheromone capture determines the olfactory performance of a pectinate antenna, which has been a challenging problem for a long time [52, 10, 28, 29]. It relies on a faithful geometry. However, because of its complexity, we had to make approximations on other aspects of the problem and restricted, for example, our analysis to steady state regime. We also modeled a filiform antenna as a perfectly smooth cylinder perpendicular to the flow. Previous studies have shown that antenna orientation [58] or presence of scales [67] could enhance their olfactory performance. It should however be noted that not all filiform antennae bear scales. We discuss the limitations of our approach in turn, before moving to the co-occurrence of designs in moths and other insects.

### Modeling a multiscale object

One of the main challenges of this work was modeling a multiscale pectinate antenna. 28000 sensilla sit on 100 rami, themselves sitting on a 10 mm long flagellum. Simulations of such structures are computer-intensive, even by today’s standards (almost two million CPU hours for the full-model simulations presented in this work). Fabrication of physical models is also exceedingly difficult, as explained in the introduction. To avoid this problem, we developed a method to model each microstructure (ramus + sensilla) by an equivalent macrostructure, whose increased diameter takes into account the influence of the sensilla on the flow. A similar approach has been used in [41] to incorporate microscopic details of setae via an equivalent cylinder. To validate this method, we ran computer-intensive simulations to determine the leakiness and drag of the full model, *i.e*. with the sensilla. We obtained similar results with both approaches, validating our equivalent macrostructure.

The main limitations of the method of equivalent cylinders are that the permeability should be uniform and the object should be flat enough to be considered as a permeable plate in order to be modeled by a row of cylinders. The method of equivalent cylinders could thus be generalized to this type of flat permeable objects in order to calculate their leakiness.

The models of pectinate antenna developed here have all a rectangular shape whereas real pectinate antennae have a leaf-like shape. In others words, the length of the rami varies along the flagellomere in a real antenna. Figure 4 gives some insight on what could happen in an antenna with varying ramus lengths. Leakiness varies, especially at high velocities (Fig4. A). However, drag and pheromone capture remain almost linear to ramus length (Fig4. B and C) and, thus, the capture-per-energy is almost independent of ramus length, except for very short rami (Fig4. D). These results were found for rectangular structures with equal-length rami and not for a leaf-shape structure but they also show that a leaf-like shape instead of a rectangular one should not change the results too much.

### Interaction between microstructures

The interactions of the rami with each other imply that combining the two levels of the antenna in an entire organ decreases the leakiness compared to the microstructure alone (one ramus and its sensilla) [30] (Fig. S11), as could be expected from fluid dynamics considerations. For velocities inferior or equal to 0.2 m s^−1^, the leakiness of an entire structure is less than 5% and the antenna behaves like a solid plate. It is known that some of the smallest insects exploit this phenomenon to decrease the weight of their wings while retaining essentially the aerodynamic properties of a membranous wing [19]. However, at high velocities, the respective leakinesses of the pectinate antenna and the microstructure tend to get closer because the boundary layers around the rami and the sensilla decrease with velocity. Compared to the microstructure, for a given air velocity, the airflow through a complete antenna is thus slowed down and molecules have more time to diffuse. The tip of the sensilla facing the flow captures then more molecules than the base of the sensilla, a phenomenon which has already been described in previous works [36, 30] (for more details see SI.)

### Co-occurrence of two designs of antennae

In terms of absolute capture, pectinate antennae always capture more molecules than filiform ones because of their larger surface. A possible explanation for maintaining a simpler filiform structure might be the aerodynamical cost of a pectinate antenna, an elevated drag. In terms of capture per unit of energy spent in the drag, filiform antennae are indeed more efficient at low velocities, below 1 m/s. For both filiform and pectinate antennae, as the air velocity increases, the drag increases faster than the pheromone capture (Figs. 3A, 2D) so the capture-per-energy (CPE) decreases in both designs (Fig. 3C). However, their respective rate of decrease does not scale identically. Although the drag scales roughly identically, it is not the case of the pheromone capture which increases faster for the pectinate antenna. As a result, CPE of the pectinate antenna scales approximately with 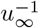 whereas in the case of the filiform antenna, it scales with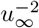 (Fig. 3C). The consequence of the different rate of decrease is that CPE of the pectinate antenna becomes superior to the one of the filiform antenna for air velocities superior to a Schmidt-number dependent threshold. For increasing Schmidt-numbers, the threshold velocity decreases. Thus, it seems that pectinate antennae should be favored in the case of high Schmidt number. However, no correlation was found between antennal shape and molecular weight of the pheromone [62]. Most pheromones have a Schmidt number around 3.5 (for more details, see the SI), a narrow range explaining maybe why no correlation was found between antennal shape and molecular weight of the pheromone [62]. The range of velocities where the CPE of pectinate antennae is superior to the one of filiform antennae corresponds roughly to the flight velocities, which range between 0.5 and 3 m/s for most species [2, 3, 30]. Thus, it seems that two designs emerged among Lepidoptera: filiform antennae for small moths moving slowly and pectinate antennae for big moths moving fast. This trade-off between filiform antenna capturing little pheromone but with little drag and pectinate antenna capturing more but at the cost of a higher drag is in accordance with the general pattern that pectinate antennae appear more often in big [62] and fast [33] moths.

Exceptions to the rule above however abound. For example, pectinate antennae do exist for small species of moths, among Geometridae for example. On the other hand, bigger moths of the Sphingidae family usually have filiform antennae. The separation between pectinate antennae on big moths and filiform antennae on small ones is thus only a trend and does not capture the whole diversity which exists in nature. Other considerations should be investigated. In the following, we treat several of them in turn according to their physical size, from large to small, spanning several orders of magnitudes.

Considering first the size of the animal, the Sphingidae appear as exemplary cases of large and very fast moths with simple filiform antennae. Interestingly, their two closely related families, the Saturniidae and the Bombycidae, have evolved pectinate antennae. So it seems that there might be a disadvantage of pectinate antennae at those very high velocities, which only Sphingidae reach (up to 15 m s^−1^ [48]). Many Sphingidae have indeed filiform antennae with small rami or, equivalently, pectinate antennae with reduced rami. At these high velocities, drag may be too important or, most likely, pectinate antennae may not support such constraint and bend. Interestingly, the antennae of the Sphingidae bear the longest sensilla trichodea ever found, up to 500 µm [37]. These long sensilla may be adaptations to increase pheromone capture at high flight velocities.

At the size of the antenna, different mechanisms might apply to the surprisingly small-sized pectinate antennae of less than half a centimeter observed in *Geometridae* [20], *Noctuidae* [24], *Nolidae* [34], Diptera [47], Hymenoptera [44]. The antenna could, for example, act as a non-leaky plate where odor reaches the surface by diffusion alone (like hermit crab appendages [66]). In this case, the pectinate structure would mainly serve to reduce the weight of the antenna.

At the size of a ramus, the relative importance of edge effects due to ramus length on the leakiness and, thus, on the olfactory performance of a pectinate antenna (Fig 4:A) has to be considered. For example, antennae with small rami have a higher leakiness than antennae with long rami at low air velocities, below m/s, for the same inter-ramus spacing. Such considerations might partially explain the relative benefits and costs of bearing pectinate antennae for small insects.

At the smallest scale, the scale of a sensillum, we observe that changing the length of the sensilla greatly influences the leakiness of the antenna and its olfactory performance (Fig. 4B). Sensillum length, or density, might then be an adaptive trait for moths to their specific ecological niche, as observed for bees as function of the degree of sociality [69]. Regarding sensillum types, trichoid and placodeum sensilla should not have the same influence on the flow. Placodeum sensilla, which are flat [23], are expected to have a lower impact on the drag and to provide a higher leakiness to the antenna.

These are only a subset of possible influences on the prevalence of filiform vs pectinate antennae. Others factors could include lifetime for example, with short lifetimes as in the non-feeding adult stages of Bombycidae and Saturnidae permitting fragile structures to exist while longer lifetimes may need more robust designs coping with repeated strong bending and dusty environment. How these different constraints work together is not obvious and will need dedicated studies. For example, our sensitivity analysis on the ramus and sensillum lengths indicate that the former tends to determine the overall aerodynamical size of the antenna and, hence, its drag, while the later impacts mainly its permeability and odor capture properties. While the filiform and pectinate designs represent the two extreme morphological types of what is in fact a continuum of designs, the lack of understanding of the ontogeny of antennal segments and their ornamentation [50] precludes us to propose a credible hypothesis for the existence of intermediate forms. Mapping the location, orientation, size and shape of all the elements of an antenna is therefore a requisite for understanding these multiscale structures.

## Conclusion

Two main architectural designs of antennae among Lepidoptera were historically studied: the spectacular but infrequent pectinate design and the more common filiform one. Their cooccurence may partly rely on their distinct mode of functioning. On one hand, filiform antennae capture few molecules but they also have a low drag. They do not cost much but they do not yield much either. On the other hand, pectinate antennae are better at capturing pheromone molecules but this improved performance is only possible at the cost of an increased drag. Thus pectinate antennae yield much but their cost is also high. Once expressed in terms of capture-per-energy, a clear pattern emerges: filiform antennae are more efficient at low velocities, whereas pectinate antennae perform better at higher ones. At which air velocity the transition occurs depends on many parameters such as Schmidt number, sensillum shape and density, inter-ramus spacing and flight velocity. More generally, considerations about the relative importance of trade-offs stemming from fluid dynamics, in particular drag, and molecule capture might explain the continuum between filiform and pectinate antennae in Lepidoptera and, more generally, the large variety of forms and architectures seen in insect antennal designs.

## References

[1] J.D. Allison and R.T. Cardé. Pheromone communication in moths: evolution, behavior, and application. University of California Press, 2016.

[2] T.C. Baker and R.G. Vogt. Measured behavioural latency in response to sex-pheromone loss in the large silk moth antheraea polyphemus. Journal of Experimental Biology, 137:29–38, 1988.

[3] J.R. Barber, B.C. Leavell, A.L. Keener, J.W. Breinholt, B.A. Chadwell, C.J.W. McClure, G.M. Hill, and A.Y. Kawahara. Moth tails divert bat attack: Evolution of acoustic deflection. Proceedings of the National Academy of Sciences, 112:2812–2816, 2015.

[4] G.A. Bartholomew and T.M. Casey. Oxygen consumption of moths during rest, pre-flight warm-up, and flight in relation to body size and wing morphology. Journal of Experimental Biology, 76:11–25, 1978.

[5] G.A. Bartholomew, D. Vleck, and C.M. Vleck. Instantaneous measurements of oxygen consumption during pre-flight warm-up and post-flight cooling in sphingid and saturniid moths. Journal of Experimental Biology, 90:17–32, 1981.

[6] Alexandra L. Basolo and Guillermina Alcaraz. The turn of the sword: length increases male swimming costs in swordtails. Proceedings of the Royal Society of London. Series B: Biological Sciences, 270:1631–1636, 8 2003.

[7] Gary J. Blomquist and R.G. Vogt, editors. Insect Pheromone Biochemistry and Molecular Biology. Academic Press, 2003.

[8] Ring T. Cardé. Navigation along windborne plumes of pheromone and resource-linked odors. Annual Review of Entomology, 66:317–336, 1 2021.

[9] A. Celani, E. Villermaux, and M. Vergassola. Odor landscapes in turbulent environments. Physical Review X, 4:1–17, 2014.

[10] A.Y.L. Cheer and M.A.R. Koehl. Fluid flow through filtering appendages of insects. Journal of Mathematics Applied in Medicine and Biology, 4:185– 199, 1987.

[11] A.Y.L. Cheer and M.A.R. Koehl. Paddles and rakes: Fluid flow through bristled appendages of small organisms. Journal of Theoretical Biology, 129:17–39, 11 1987.

[12] Christopher James Clark and Robert Dudley. Flight costs of long, sexually selected tails in hummingbirds. Proceedings of the Royal Society B: Biological Sciences, 276:2109–2115, 6 2009.

[13] C. Cummins, M. Seale, A. Macente, D. Certini, E. Mastropaolo, I.M. Viola, and N. Nakayama. A separated vortex ring underlies the flight of the dandelion. Nature, 562:414–418, 2018.

[14] Mark A. Elgar, Tamara L. Johnson, and Matthew R.E. Symonds. Sexual selection and organs of sense: Darwin’s neglected insight. Animal Biology, 69:63–82, 2019.

[15] Mark A. Elgar, Dong Zhang, Qike Wang, Bernadette Wittwer, Hieu Thi Pham, Tamara L Johnson, Christopher B Freelance, and Marianne Coquilleau. Insect antennal morphology: The evolution of diverse solutions to odorant perception. The Yale journal of biology and medicine, 91:457–469, 12 2018.

[16] T. Engels, K. Schneider, J. Reiss, and M. Fagre. A wavelet-adaptive method for multiscale simulation of turbulent flows in flying insects. Communications in Computational Physics, 30:1118–1149, 6 2021.

[17] Thomas Engels, Dmitry Kolomenskiy, and Fritz-Olaf Lehmann. Flight efficiency is a key to diverse wing morphologies in small insects. Journal of The Royal Society Interface, 18, 10 2021.

[18] Thomas Engels, Hung Truong, Marie Farge, Dmitry Kolomenskiy, and Kai Schneider. Computational aerodynamics of insect flight using volume penalization. Comptes Rendus. Mécanique, 350:1–20, 11 2022.

[19] S. Farisenkov, D. Kolomenskiy, P. Petrov, N. Lapina, T. Engels, F.-O. Lehmann, R. Onishi, H. Liu, and A. Polilov. Novel flight style and light wings boost flight performance of tiny beetles. Nature, 2021.

[20] W. T. M. Forbes. Pectinate antennae in the geometridae (lepidoptera). Psyche: A Journal of Entomology, 32:106–112, 1 1925.

[21] Taiping Gao, Chungkun Shih, Conrad C. Labandeira, Jorge A. Santiago-Blay, Yunzhi Yao, and Dong Ren. Convergent evolution of ramified antennae in insect lineages from the early cretaceous of northeastern china. Proceedings of the Royal Society B: Biological Sciences, 283:20161448, 9 2016.

[22] Jana Goyens, Sam Van Wassenbergh, Joris Dirckx, and Peter Aerts. Cost of flight and the evolution of stag beetle weaponry. Journal of The Royal Society Interface, 12:20150222, 5 2015.

[23] Eric Hallberg and Bill S. Hansson. Arthropod sensilla: Morphology and phylogenetic considerations. Microscopy Research and Technique, 47:428–439, 12 1999.

[24] B.S. Hansson, S. Anton, and T.A. Christensen. Structure and function of antennal lobe neurons in the male turnip moth, agrotis segetum (lepidoptera: Noctuidae). Journal of Comparative Physiology A, 175, 11 1994.

[25] Bernd Heinrich and Timothy M. Casey. Metabolic rate and endothermy in sphinx moths. Journal of Comparative Physiology, 82:195–206, 1973.

[26] Francisco J.H. Heras and Simon B. Laughlin. Investments in photoreceptors compete with investments in optics to determine eye design. BioRxiv Preprint, January, 2024.

[27] J. Howard, B. Blakeslee, and S.B. Laughlin. The intracellular pupil mechanism and photoreceptor signal: noise ratios in the fly lucilia cuprina. Proceedings of the Royal Society of London. Series B. Biological Sciences, 231:415–435, 9 1987.

[28] J.A.C. Humphrey and H. Haj-Hariri. Detection and real time processing of odor plume information by arthropods in air and water. In Proceedings of the AFI-2002 Mini-Symposium on Advanced Fluids Information—Fusion of EFD and CFD, pages 47–67, 2002.

[29] M. Jaffar-Bandjee, G.J.M. Krijnen, and J. Casas. Challenges in modeling pheromone capture by pectinate antennae. Integrative and Comparative Biology, page icaa057, 2020.

[30] M. Jaffar-Bandjee, T. Steinmann, G.J.M. Krijnen, and J. Casas. Insect pectinate antennae maximize odor capture at intermediate flight speeds. Proceedings of the National Academy of Sciences, page 202007871, 2020.

[31] M. Jaffar-Bandjee, T. Steinmann, G.J.M. Krijnen, and J. Casas. Leakiness and flow capture ratio of insect pectinate antennae. Journal of the Royal Society Interface, 17:20190779, 2020.

[32] Ludovic Jami, Thomas Zemb, Jérôme Casas, and Jean François Dufrêche. How adsorption of pheromones on aerosols controls their transport. ACS Central Science, 6:1628–1638, 2020.

[33] Tamara L. Johnson, Mark A. Elgar, and Matthew R. E. Symonds. Movement and olfactory signals: Sexually dimorphic antennae and female flight-lessness in moths. Frontiers in Ecology and Evolution, 10, 8 2022.

[34] Tamara L. Johnson, Matthew R. E. Symonds, and Mark A. Elgar. Sexual selection on receptor organ traits: younger females attract males with longer antennae. The Science of Nature, 104:44, 6 2017.

[35] Nirag Kadakia, Mahmut Demir, Brenden T. Michaelis, Brian D. DeAngelis, Matthew A. Reidenbach, Damon A. Clark, and Thierry Emonet. Odour motion sensing enhances navigation of complex plumes. Nature, 611:754–761, 11 2022.

[36] S. Kanaujia and K.-E. Kaissling. Interactions of pheromone with moth antennae: Adsorption, desorption and transport. Journal of Insect Physiology, 31:71–81, 1985.

[37] Thomas A. Keil. Fine structure of the pheromone-sensitive sensilla on the antenna of the hawkmoth, manduca sexta. Tissue and Cell, 21:139–151, 1 1989.

[38] Thomas A. Keil and R. Alexander Steinbrecht. Mechanosensitive and Olfactory Sensilla of Insects, pages 477–516. Springer US, 1984.

[39] Bo-Youn Kim, Young-Whan Park, Nam-Sook Park, and Sang-Mong Lee. Collection and characteristics of the wild silkmoth, samia cynthia pryeri, in korea. International Journal of Industrial Entomology, 3:101–103, 2001.

[40] Darius Klassen, Martin D. Lennox, Marie-Josée Dumont, Gérald Chouinard, and Jason R. Tavares. Dispensers for pheromonal pest control. Journal of Environmental Management, 325:116590, 1 2023.

[41] Dmitry Kolomenskiy, Sergey Farisenkov, Thomas Engels, Nadezhda Lapina, Pyotr Petrov, Fritz-Olaf Lehmann, Ryo Onishi, Hao Liu, and Alexey Polilov. Aerodynamic performance of a bristled wing of a very small insect. Experiments in Fluids, 61:194, 9 2020.

[42] H. Kramers. Heat transfer from spheres to flowing media. Physica, 12:61–80, 1946.

[43] Niels P. Kristensen. 4. Skeleton and muscles: adults, pages 39–132. De Gruyter, 12 2003.

[44] S.Y. Li. Notes on larval instars and adult antennae of neodiprion abietis (hymenoptera: Diprionidae). The Canadian Entomologist, 135:745–748, 10 2003.

[45] C. Loudon and M.A.R. Koehl. Sniffing by a silkworm moth: wing fanning enhances air penetration through and pheromone interception by antennae. Journal of Experimental Biology, 203:2977–2990, 2000.

[46] Catherine Loudon. Antennae, pages 21–23. Academic Press, second edition, 2009.

[47] Loïc Matile. A new australian genus of keroplatidae with pectinate antennae (diptera: Mycetophiloidea). Australian Journal of Entomology, 20:207–212, 8 1981.

[48] Keith C. McKeown. Insect wonders of Australia. Angus & Robertson, 1935.

[49] Aryeh H. Miller, James T. Stroud, and Jonathan B. Losos. The ecology and evolution of key innovations. Trends in Ecology & Evolution, 38:122–131, 2 2023.

[50] A. Minelli. The insect antenna: segmentation, patterning and positional homology. Journal of Entomological and Acarological Research, 49, 4 2017.

[51] J. Murlis, J.S. Elkinton, and R.T. Cardé. Odor plumes and how insects use them. Annual Review of Entomology, 37:505–532, 1992.

[52] J.D. Murray. Reduction of dimensionability in diffusion processes: antenna receptors of moths, pages 83–127. Oxford University Press, 1977.

[53] J.K. Nayar and E. van Handel. Flight performance and metabolism of the moth spodoptera frugiperda. Journal of Insect Physiology, 17:2475–2479, 1971.

[54] Stefanie Neupert, Graham A. McCulloch, Brodie J. Foster, Jonathan M. Waters, and Paul Szyszka. Reduced olfactory acuity in recently flightless insects suggests rapid regressive evolution. BMC Ecology and Evolution, 22:50, 12 2022.

[55] Jeremy E. Niven and Simon B. Laughlin. Energy limitation as a selective pressure on the evolution of sensory systems. Journal of Experimental Biology, 211:1792–1804, 6 2008.

[56] Hieu T. Pham, Mark A. Elgar, Emile van Lieshout, and Kathryn B. McNamara. Experimental immune challenges reduce the quality of male antennae and female pheromone output. Scientific Reports, 12:3578, 2022.

[57] Yangang Ren, Max R. McGillen, Véronique Daële, Jérôme Casas, and Abdelwahid Mellouki. The fate of methyl salicylate in the environment and its role as signal in multitrophic interactions. Science of the Total Environment, 749:141406, 2020.

[58] Thomas L. Spencer, Nina Mohebbi, Guangyuan Jin, Matthew L. Forister, Alexander Alexeev, and David L. Hu. Moth-inspired methods for particle capture on a cylinder. Journal of Fluid Mechanics, 884:A34, 2 2020.

[59] R.A. Steinbrecht. Zur morphometrie der antenne des seidenspinners, bombyx mori l.: Zahl und verteilung der riechsensillen (insecta, lepidoptera). Zeitschrift für Morphologie der Tiere, 68:93–126, 1970.

[60] R. D. Stevenson and R. K. Josephson. Effects of operating frequency and temperature on mechanical power output from moth flight muscle. Journal of Experimental Biology, 149:61–78, 1990.

[61] R.K. Suarez. Energy metabolism during insect flight: Biochemical design and physiological performance. Physiological and Biochemical Zoology, 73:765–771, 2000.

[62] M.R.E. Symonds, T.L. Johnson, and M.A. Elgar. Pheromone production, male abundance, body size, and the evolution of elaborate antennae in moths. Ecology and Evolution, 2:227–246, 2011.

[63] Paul Szyszka, Thierry Emonet, and Timothy L Edwards. Extracting spatial information from temporal odor patterns: insights from insects. Current Opinion in Insect Science, 59:101082, 10 2023.

[64] M. J. Tang, M. Shiraiwa, U. Pöschl, R. A. Cox, and M. Kalberer. Compilation and evaluation of gas phase diffusion coefficients of reactive trace gases in the atmosphere: Volume 2. diffusivities of organic compounds, pressure-normalised mean free paths, and average knudsen numbers for gas uptake calculations. Atmospheric Chemistry and Physics, 15:5585–5598, 2015.

[65] Steven Vogel. How much air passes through a silkmoth’s antenna? Journal of Insect Physiology, 29:597–602, 1 1983.

[66] L.D. Waldrop and M.A.R. Koehl. Do terrestrial hermit crabs sniff? air flow and odorant capture by flicking antennules. Journal of the Royal Society, Interface, 13:20150850, 1 2016.

[67] Q. Wang, Y. Shang, D.S. Hilton, K. Inthavong, D. Zhang, and M.A. Elgar. Antennal scales improve signal detection efficiency in moths. Proceedings of the Royal Society B, 285:20172832, 2018.

[68] C H K Williamson. Vortex dynamics in the cylinder wake. Annual Review of Fluid Mechanics, 28:477–539, 1 1996.

[69] Bernadette Wittwer, Abraham Hefetz, Tovit Simon, Li E. K. Murphy, Mark A. Elgar, Naomi E. Pierce, and Sarah D. Kocher. Solitary bees reduce investment in communication compared with their social relatives. Proceedings of the National Academy of Sciences, 114:6569–6574, 6 2017.

