## Supplemental Information for "Olfactory performance explains duality of antennal architectural designs in Lepidoptera"

#### Diffusion Coefficients of Sex Pheromones

We selected several species of moths and calculated an approximate estimation of their respective sex pheromone. The objective was to obtain a range of diffusion coefficients $\nu_{\text{phero}}$ in air of sex pheromones. We estimated the diffusion coefficients based on the formula based on the chemical formula of the pheromones [23]. From those data, we define the range of $\nu_{\text{phero}} \in \left[10^{-6}, 10^{-5}\right]$, yielding the range of Schmidt numbers used in the main text.

| Type I pheromone | | | | |
|---|---|---|---|---|
| Diffusion coefficient ($m^2\,s^{-1}$) | Chemical formula | Moth species | Family | Reference |
| $4.66 \times 10^{-6}$ | $C_{16}H_{30}O$ | *Bombyx mori* | Bombycidae | [9] |
| $4.80 \times 10^{-6}$ | $C_{14}H_{27}O_3N$ | *Bucculatrix thurberiella* | Bucculatricidae | [13] |
| $4.83 \times 10^{-6}$ | $C_{13}H_{25}O_3N$ | *Bucculatrix thurberiella* | Bucculatricidae | [13] |
| $4.70 \times 10^{-6}$ | $C_{16}H_{28}O$ | *Desmia funeralis* | Crambidae | [20] |
| $3.62 \times 10^{-6}$ | $C_{26}H_{44}O_2$ | *Euproctis chrysorrhoea* | Erebidae | [18] |
| $3.77 \times 10^{-6}$ | $C_{24}H_{42}O_2$ | *Euproctis pulverea* | Erebidae | [28] |
| $3.61 \times 10^{-6}$ | $C_{26}H_{46}O_2$ | *Euproctis pulverea* | Erebidae | [28] |
| $3.82 \times 10^{-6}$ | $C_{23}H_{44}O_2$ | *Euproctis similis* | Erebidae | [31] |
| $4.97 \times 10^{-6}$ | $C_{14}H_{24}O_2$ | *Phyllonorycter mespilella* | Gracillariidae | [10] |
| $4.66 \times 10^{-6}$ | $C_{16}H_{30}O$ | *Heliothis virescens* | Noctuidae | [22] |
| $4.93 \times 10^{-6}$ | $C_{14}H_{26}O_2$ | *Trichopulsia ni* | Noctuidae | [8] |
| $4.39 \times 10^{-6}$ | $C_{18}H_{30}O_2$ | *Thaumetopoea processionea* | Notodontidae | [12] |
| $4.36 \times 10^{-6}$ | $C_{18}H_{32}O_2$ | *Thaumetopoea processionea* | Notodontidae | [12] |
| $4.76 \times 10^{-6}$ | $C_{15}H_{28}O_2$ | *Nudaurelia cytherea cytherea* | Saturniidae | [14] |

Table 1: Diffusion coefficients $\nu_{\mathrm{phero}}$ in air of several Type I sex pheromones of moths [1].

| Type II pheromone | | | |
|---|---|---|---|
| Diffusion coefficient ($m^2 s^{-1}$) | Chemical formula | Family | Reference |
| $4.30 \times 10^{-6}$ | $C_{19}H_{36}$ | Geometridae | [1] |
| $4.19 \times 10^{-6}$ | $C_{20}H_{38}$ | Arctiidae Geometridae | [1] |
| $4.08 \times 10^{-6}$ | $C_{21}H_{40}$ | Arctiidae Geometridae Noctuidae | [1] |
| $4.38 \times 10^{-6}$ | $C_{18}H_{34}O$ | Geometridae | [1] |
| $4.26 \times 10^{-6}$ | $C_{19}H_{36}O$ | Geometridae | [1] |
| $4.04 \times 10^{-6}$ | $C_{21}H_{40}O$ | Arctiidae Noctuidae | [1] |
| $4.60 \times 10^{-6}$ | $C_{17}H_{30}$ | Geometridae | [1] |
| $4.46 \times 10^{-6}$ | $C_{18}H_{32}$ | Geometridae | [1] |
| $4.33 \times 10^{-6}$ | $C_{19}H_{34}$ | Geometridae | [1] |
| $4.21 \times 10^{-6}$ | $C_{20}H_{36}$ | Geometridae Noctuidae | [1] |
| $4.10 \times 10^{-6}$ | $C_{21}H_{38}$ | Geometridae Lymantriidae Noctuidae | [1] |
| $3.90 \times 10^{-6}$ | $C_{23}H_{42}$ | Arctuidae Pyralidae | [1] |
| $4.55 \times 10^{-6}$ | $C_{17}H_{30}O$ | Geometridae | [1] |
| $4.41 \times 10^{-6}$ | $C_{18}H_{32}O$ | Geometridae | [1] |
| $4.29 \times 10^{-6}$ | $C_{19}H_{34}O$ | Geometridae Lymantriidae | [1] |
| $4.17 \times 10^{-6}$ | $C_{20}H_{36}O$ | Arctiidae Noctuidae | [1] |
| $4.06 \times 10^{-6}$ | $C_{21}H_{38}O$ | Arctiidae Lymantriidae Noctuidae | [1] |
| $4.36 \times 10^{-6}$ | $C_{19}H_{32}$ | Geometridae | [21, 26, 29] |
| $4.12 \times 10^{-6}$ | $C_{21}H_{36}$ | Geometridae | [32] |
| $4.21 \times 10^{-6}$ | $C_{20}H_{33}O$ | Arctiidae | [24] |
| $4.10 \times 10^{-6}$ | $C_{21}H_{35}O$ | Arctiidae | [24] |
| $4.02 \times 10^{-6}$ | $C_{21}H_{38}O_2$ | Lymantriidae | [11, 30] |

Table 2: Diffusion coefficients $\nu_{\text{phero}}$ in air of several Type II sex pheromones of moths [1].

| Type III pheromone | | | | |
|---|---|---|---|---|
| Diffusion coefficient ($m^2\,s^{-1}$) | Chemical formula | Insect species | Family | Reference |
| $7.01 \times 10^{-6}$ | $C_7H_{14}O$ | *Eriocrania cicatricella* | Eriocraniidae | [32] |
| $7.00 \times 10^{-6}$ | $C_7H_{16}O$ | *Eriocrania cicatricella* | Eriocraniidae | [32] |
| $6.31 \times 10^{-6}$ | $C_9H_{16}O$ | *Eriocrania sangii*<br>*Eriocrania semipurpurella* | Eriocraniidae | [16] |
| $6.40 \times 10^{-6}$ | $C_9H_{14}O$ | *Stigmella malella* | Nepticulidae | [25] |
| $4.86 \times 10^{-6}$ | $C_{14}H_{30}O_2$ | *Thyridopteryx ephemeraeformis*<br>*Eriocrania semipurpurella* | Eriocraniidae | [16] |

Table 3: Diffusion coefficients $\nu_{\text{phero}}$ in air of several type III sex pheromones of moths [1].

### Filiform antenna

A drag force $F_\mathrm{d}$ is created when a filiform antenna encounters an airflow, which is important from an energetic point of view since animals carry the antenna with them as they fly. It can be computed as

$$F_\mathrm{d} = C_\mathrm{d}\frac{1}{2}\varrho_\mathrm{air}u_\infty^2 A_\mathrm{fili} \tag{1}$$

where $C_\mathrm{d}$ is the drag coefficient, $\varrho_\mathrm{air}$ the density of air, $u_\infty$ the cruising velocity and $A_\mathrm{fili} = L_\mathrm{fili}d_\mathrm{fili}$ the projected area. The drag coefficient depends on the Reynolds number $\mathrm{Re} = u_\infty d/\nu_\mathrm{air}$. Depending on the exact Reynolds number, many empirical formula exist to calculate $C_\mathrm{d}$. Here, the empirical formula proposed in [19] was chosen because it is valid for a large range of Reynolds numbers,

$$C_\mathrm{d} = 1.38\mathrm{Re}^{-0.05} + 7.72\mathrm{Re}^{-0.69} + 1.82\mathrm{Re}^{-1}. \tag{2}$$

We also run 2D finite element (FEM) simulations with the commercial software Comsol (v5.6) to determine the olfactory performance of filiform antennae, modeled as smooth cylinders. We test two diameters: 60 and 120 µm. To compare with the pectinate antenna, we assume that the filiform antennae are 1 cm long. Thus, the aspect ratio (length over diameter) is over 100. As a consequence, we assume that edge effects would have little influence on the performance of the entire antenna and reduce the problem to a 2D simulation. In case of absolute values such as drag and absolute capture of pheromone, we multiply the results by the length of the antenna. The simulation domain is equal to 100 × 100 mm and the cylinder is located its center.

To simulate the flow of air, we use the *Laminar flow* package of Comsol (v5.6). We resort to the *Transport of Diluted Species* for the pheromone concentration and use the multiphysics tool *Reacting flow, Diluted species* to link fluid dynamics and mass transport. The model is solved in a steady state using the PARDISO solver. There is a no-slip boundary condition at the surface of the cylinder. The normal velocity is fixed equal to $u_\infty$ on the upstream side and a zero pressure condition is set on the downstream side. The two remaining sides (top/bottom) are walls moving at $u_\infty$ tangentially to the flow, *i.e.* in the $x$ direction (Fig. 1). Regarding mass transfer, the concentration is zero at the surface of the cylinders and fixed at $1\,\mathrm{mol\,m^{-3}}$ on the upstream side. On the downstream side, the gradient of the concentration with respect to the normal of the downstream side, *i.e.* $\frac{\partial c}{\partial x}$, is fixed zero. The two remaining sides have a no flux condition (Fig. 1).

The mesh has 27 894 elements. They are all triangular except close to the cylinders where three levels of boundary layers are modeled with quadrilateral elements. The minimum and maximum sizes of mesh elements are fixed at 15 µm and 1.3 mm, respectively. Close to the cylinder, we refine the mesh using a square of size 2 × 2 mm centered on the cylinder. Inside this square, the mesh is refined, and minimum and maximum element sizes are 2 µm and 670 µm, respectively.

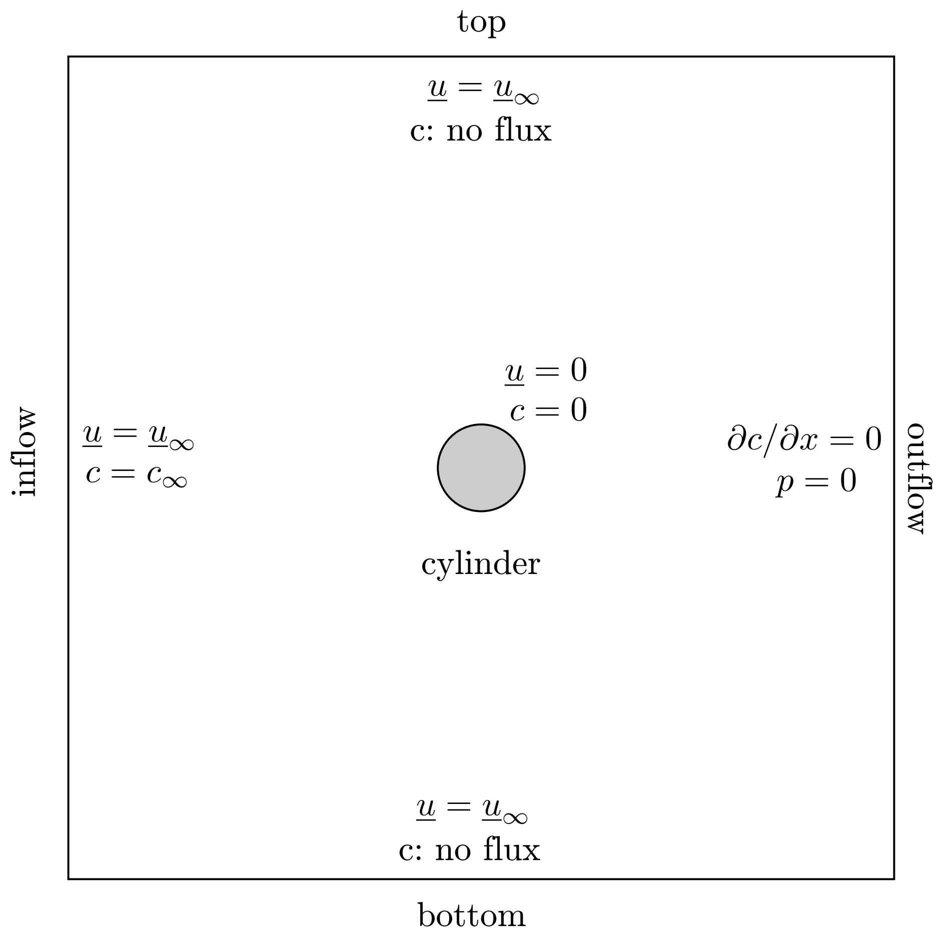


Figure 1: Simulation domain and boundary conditions in the case of a filiform antenna. In order to increase the clarity, the picture is not to scale.

#### Simulations to determine the equivalent macrostructures and the pheromone capture by the sensilla

Determining the diameter of the equivalent macrostructure requires two steps: (a) simulation of the microstructure to determine its drag at a given velocity and (b) from the resulting drag determination of the corresponding diameter for equal drag.

For step (a), we perform a simulation with the setup shown in Fig. 2. The domain borders in the $z$ and $x$ direction are periodic; only the $y$ direction which corresponds to the inflow direction is nonperiodic. Such a simulation is thus equivalent to a microstructure of infinite length in $z$ direction, embedded in an infinite array of identical microstructures in the $x$ direction. Those simulations yield the drag force $F_d$ of the microstructure for all values of $u_\infty$.

For step (b), we perform simulations using the same setup but without the sensilla (Fig. 3). For each value of $u_\infty$, we sample a number of different $d_{\text{cyl}}$ and determine the drag. We plot the resulting $F_d$ as a function of $d_{\text{cyl}}$ (Fig. 4) and determine the value of $\widetilde{d}_{\text{rami}}$ for equal drag (Table 4).

We use the same model as in step (a) to determine the pheromone capture by the sensilla of a pectinate antenna. We add the mass transport equation to the simulation. The far-field concentration is set at $1\,\text{mol}\,\text{m}^{-3}$ and the concentration at the surface of the sensilla is assumed null.

This model only considers capture of pheromone within the air flowing through the antenna — we exclude diffusion of pheromone from the air passing around it. This assumption is correct if the pheromone diffusing from the air passing around the antenna is negligible compared to the pheromone captured in the flux going through it. This is however not the case at low velocities: due to the very low Le at low $u_\infty$, most of the air and pheromone is deflected around the antenna and the air flowing between the rami and the sensilla is very slow. To assess the importance of this phenomenon, we calculate the Péclet number, which compares the importance of transport by advection and diffusion:

$$\text{Pe} = \frac{v_\infty \text{Le} L_{\text{rami}}}{\nu_{\text{phero}}} \tag{3}$$

At $v_\infty = 0.1\,\text{m}\,\text{s}^{-1}$, we obtain a Péclet number of 0.7. This value is close to 1 meaning both advection and diffusion are important and should be taken into account.

To correct for this modeling error at low $u_\infty$, we use the *filiform* antenna with $d_{\text{fili}} = 60\,\mu\text{m}$ as indicator. If it captures not only pheromone from the airflow facing it but also from the airflow on the sides (which is a clue that diffusion is non-negligible), its capture rate is higher than the pheromone flux in the free stream passing through the filiform antennas projected area: $\Phi_{\text{fili}} > c_\infty u_\infty A$. The corresponding excess in $\Phi$ is then added to the pheromone capture model of the pectinate antenna. This additional pheromone capture $\Phi_{\text{add}}$ is thus calculated as:

$$\Phi_{\text{add}} = \Phi_{\text{fili}} - c_\infty u_\infty A. \tag{4}$$

To validate this corrective step, we run a Comsol simulation on an equivalent macrostructure with $\widetilde{d}_{\text{rami}} = 103\,\mu\text{m}$, $u_\infty = 0.1\,\text{m}\,\text{s}^{-1}$ and include pheromone transport. If we only consider the airflow going through the antenna, the pheromone capture rate is equal to $1.5 \times 10^{-8}\,\text{mol}\,\text{s}^{-1}$ whereas the total value amounts to $11.8 \times 10^{-8}\,\text{mol}\,\text{s}^{-1}$. If we correct with the additional mass capture $\Phi_{\text{add}}$, equal to $6.6 \times 10^{-8}\,\text{mol}\,\text{s}^{-1}$ at this far-field velocity, the total mass capture of the pectinate antenna reaches $8.1 \times 10^{-8}\,\text{mol}\,\text{s}^{-1}$, which is closer to the total value calculated in the simulation.

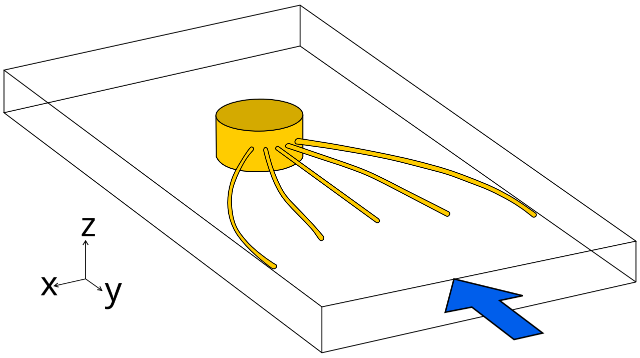


Figure 2: Simulations to determine the drag and the pheromone capture of a microstructure. 3D view of one and its five sensilla. The blue arrow shows the direction of the flow. Because rami have periodic rows of sensilla, we restrict the simulation domain to one row of five sensilla. One ramus is 25 µm high and has a diameter of 50 µm.

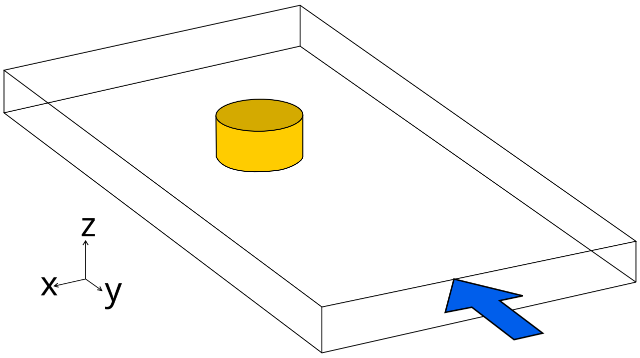


Figure 3: Simulations to determine the drag of a rami with variable diameter.

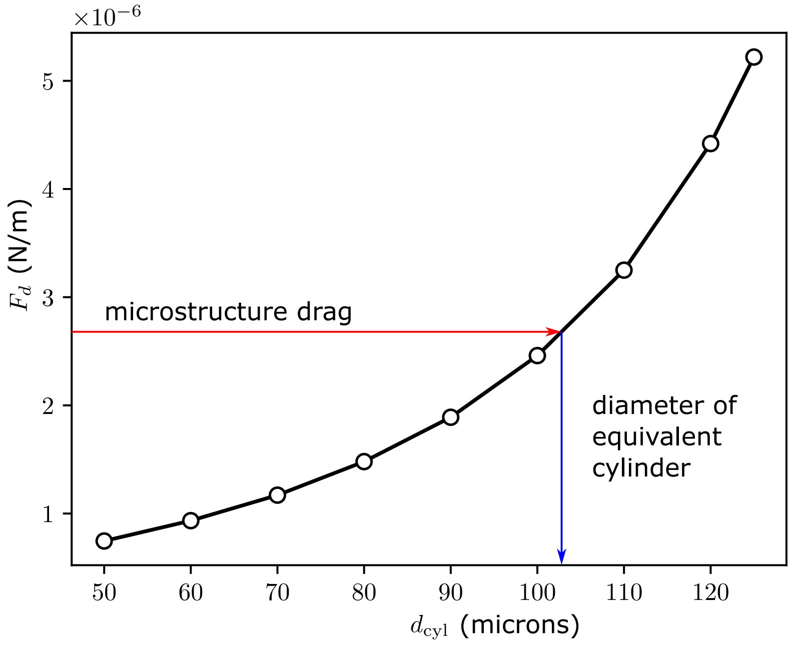


Figure 4: Determination of the diameter of the equivalent cylinder to the microstructure. The drag per cylinder and per unit of length is determined at each velocity for several diameters (black dots). The values between two data points are linearly interpolated (black line). Here, the air velocity is equal to $0.5\,\mathrm{m\,s^{-1}}$. The drag of the microstructure at this given velocity (red line) is equal to $2.7\,\mathrm{N\,mm^{-2}}$ and is used to determine the diameter of the corresponding equivalent cylinder (blue line), here approximately equal to $103\,\mu\mathrm{m}$.

| $u_\infty$ (m/s) | 0.01 | 0.02 | 0.05 | 0.1 | 0.2 | 0.5 | 1 | 2 |
|---|---|---|---|---|---|---|---|---|
| $\widetilde{d}_{\text{rami}}$ (µm) | 104.1 | 104.1 | 104.1 | 104 | 104 | 103.7 | 102.8 | 100.4 |

Table 4: Resulting equivalent diameter as a function of the inflow velocity.

#### Simulations of the equivalent macrostructure

We conduct Finite Element Method simulations (Comsol v5.6) to determine the leakiness of the equivalent macrostructure, *i.e.* a flagellum and its rami. To account for the sensilla, the rami of these macrostructures have enlarged diameters as described in the main text.

In the simulations, the macrostructures are composed of one main cylinder modeling the flagellum and 100 identical secondary cylinders modeling the rami. The flagellum is modeled by a cylinder of length 10 mm with a diameter of 250 µm. To avoid sharp edges, a sphere of diameter 250 µm is added at the two tips of the cylinder. Rami are identical cylinders of length 1.5 mm and diameter $\widetilde{d}_{\mathrm{rami}}$. Similarly to the flagellum, a sphere with the same diameter as the rami is added at their tips to avoid sharp edges. The rami are all perpendicular to the flagellum and to the flow, and are evenly distributed, 50 on each side of the flagellum, with an inter-rami distance (center-to-center) of 200 µm.

We use the Laminar flow package of Comsol (v5.6) in a steady-state regime with the GMRES (Generalized Minimal RESidual) steady-state solver. We set a no-slip boundary condition at the surface of the macrostructure. We also use two symmetric boundary conditions to reduce the model to one quarter (Fig. 5). Air velocity is set at the desired far-field velocity on the upstream boundary. Pressure is fixed at zero on the downstream boundary. The two remaining ones are defined as no-slip walls moving at the same speed and direction as the far-field velocity. We choose a simulation domain equal to $80 \times 40 \times 18$ mm. We ensured that the simulations give results close to open-field conditions, *i.e.*, the domain is big enough to have no significant influence.

Given the number and length of rami, we determine the computational mesh with special care. Subdomains around the macrostructure are defined to provide a finer mesh close to the rami (Fig. 5). The first subdomain is $5 \times 10 \times 4.5$ mm and centered on the rami. Here, the mesh is the finest with minimum element size of 10 µm and maximum element size of 100 µm. The second subdomain is $2.1 \times 6 \times 1.8$ mm with minimum element size of 100 µm and maximum element size of 1 mm. The remaining domain is meshed with minimum element size of 270 µm and maximum element size of 1.4 mm. Eventually, the mesh contains around 4 870 000 elements.

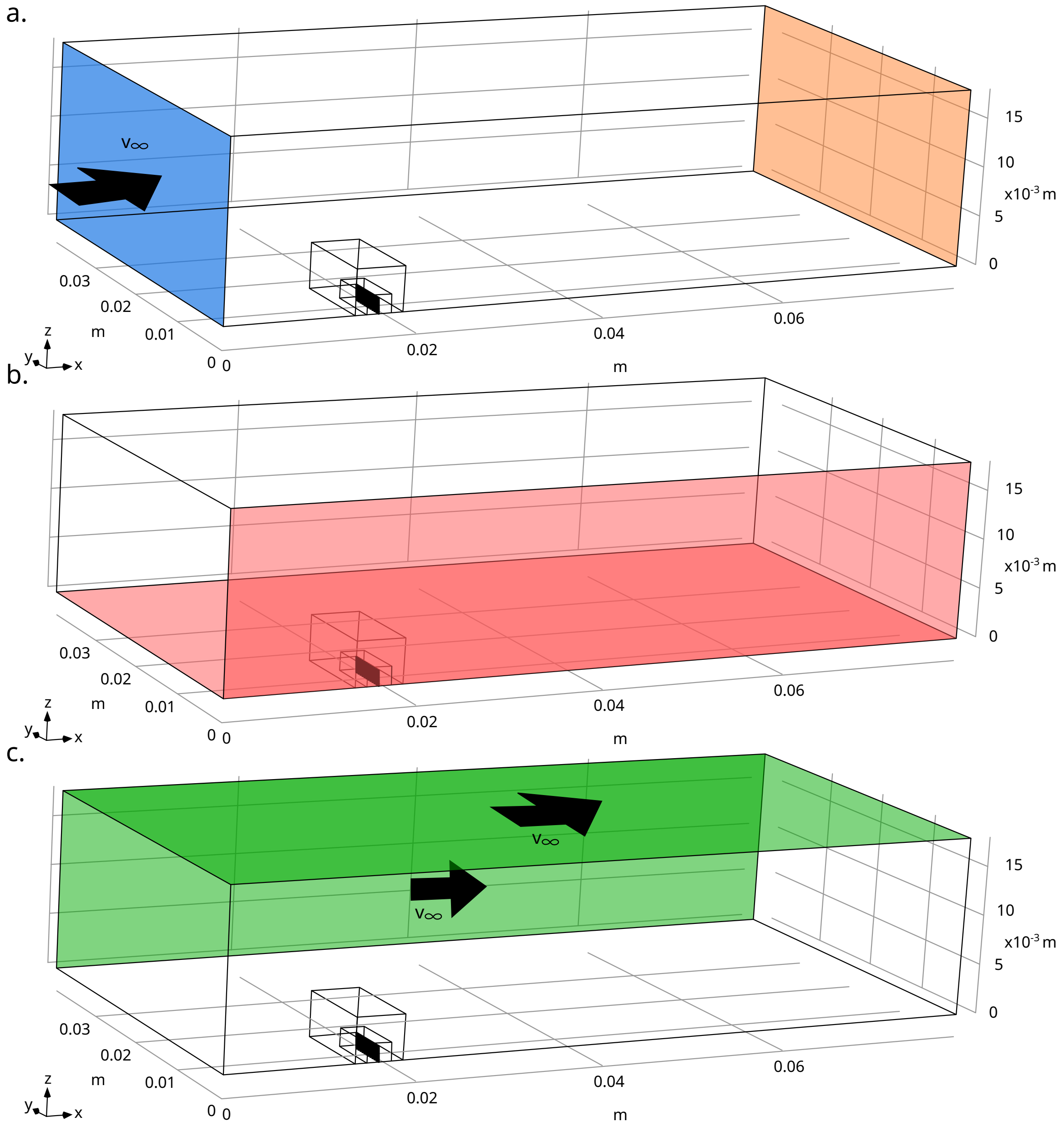


Figure 5: Simulation domain and boundary conditions in the case of a pectinate antenna. The three black boxes show the simulation domain and the two subdomains. a: The blue face is the inlet face where the normal velocity is set at $u_\infty$. The orange face is the outlet face with a pression equal to 0. b: The two red faces represent the two symmetrical boundary condition, allowing to simulate only a quarter of the entire structure. c: The green faces show the moving walls with a no-slip condition, moving at $u_\infty$ in the $x$ direction.

### Details of PIV experiments

The printed equivalent macrostructures are scaled up 10-fold, and fabricated with an Eden 260 3D printer (Stratasys, Eden Prairie, MN, USA) in VeroClear material (Stratasys, Eden Prairie, MN, USA). Particle Image Velocimetry (PIV) is used to measure the velocity fields around the structures. In PIV, the fluid of interest is seeded with particles and a laser sheet illuminates the particles in a given plane of interest. A camera records the movements of the particles and the velocity field is numerically reconstructed.

Because the 3D-printed structures are scaled up, it is more convenient to PIV measurements in other fluids, namely water and rapeseed oil. To ensure similar dynamics, the velocities in water and oil are calculated so that their Reynolds number is equal to the Reynolds number of a real antenna in air for the corresponding velocity. The Reynolds number is defined as:

$$\mathrm{Re} = \frac{du}{\nu}$$

With $d$ the characteristic length, $u$ the characteristic velocity and $\nu$ the viscosity of the fluid.

Here, the models are scaled-up by a factor $\alpha = 10$ so $d_\mathrm{f} = \alpha d_\mathrm{air}$ with $d_\mathrm{f}$ the characteristic length in the fluid (water or oil) and $d_\mathrm{air}$ the characteristic length in air.

Equal Reynolds numbers give:

$$u_\mathrm{f} = \frac{\nu_\mathrm{f}}{\nu_\mathrm{air}} \frac{1}{\alpha} u_\mathrm{air}$$

With $u_\mathrm{f}$ the velocity in water or oil, $\nu_\mathrm{f}$ the viscosity in water ($1 \times 10^{-6}\,\mathrm{m^2\,s^{-1}}$) or oil ($50 \times 10^{-6}\,\mathrm{m^2\,s^{-1}}$ at $29\,^\circ\mathrm{C}$) and $u_\mathrm{air}$ the velocity in air. The structures are immersed in a $25 \times 30 \times 50\,\mathrm{mm}$ tank either filled with water or oil depending on the investigated air velocity (Fig.6) and seeded with hollow glass particles covered with silver (10 µm; SUGS-10; Dantec Dynamics, Skovlunde, Denmark). Images are recorded perpendicular to the tank. The camera (Phantom 9.1; Vision Research, Perth, Australia) and the laser (MGL-F; 532nm; 2W CNI, Changchun, China) remain still while the tank is moved at the required velocity with a motorized linear axis LST1000D-T3, Zaber Technologies, Vancouver, British Columbia, Canada). A cover is added at the free surface to prevent wave formation except for a slit used by the support bearing the structure.

In the velocity fields obtained from the PIV measurements, the antenna obscures particles between rami. As a consequence, we can not measure directly the flow at the location of the antenna [2]. We thus performed measurements in several planes up- and downstream of the macrostructure and determined Le using the principle of continuity. To validate this method, we applied it to velocity fields obtained in the numerical simulations, where we can readily calculate the airflow through the antenna. The airflow upstream of the antenna is decreased greatly close to the macrostructure whereas the airflow downstream of the antenna varies less. Thus, we used the airflow downstream of the antenna to determine, through continuity, the flow at the location of the antenna.

To calculate the drag from the PIV data, we proceed as follows. PIV gives velocity fields and we extract the horizontal and vertical components of the velocity as well as their spatial derivatives. We resorted to the *Pressure from PIV* package from Lavision (Lavision GmbH, Göttingen, Germany) to obtain the pressure field $p$. Following [17], we then apply a momentum balance in the horizontal direction, along $\overrightarrow{e_x}$, on a volume control around the macrostructures to determine $F_\mathrm{d}$. We define

$$\vec{u} = \begin{pmatrix} u \\ v \end{pmatrix} \tag{5}$$

$$\overline{\overline{\tau}} = \begin{pmatrix} \tau_{xx} & \tau_{xy} \\ \tau_{yx} & \tau_{yy} \end{pmatrix} \tag{6}$$

where

$$\tau_{xx} = 2\mu \frac{\partial u}{\partial x} \tag{7}$$

$$\tau_{yy} = 2\mu \frac{\partial v}{\partial y} \tag{8}$$

$$\tau_{xy} = \mu \left( \frac{\partial u}{\partial y} + \frac{\partial v}{\partial x} \right) \tag{9}$$

In a steady-state, the time derivatives of the velocity vanish and the drag force is given as:

$$F_{\mathrm{d}} = -\varrho \overrightarrow{e_x} \iint_S (\vec{u} \cdot \vec{n})\, \vec{v}\, \mathrm{d}S - \overrightarrow{e_x} \iint_S p\vec{n}\, \mathrm{d}S + \overrightarrow{e_x} \iint_S \overline{\overline{\tau}}\vec{n}\, \mathrm{d}S \tag{10}$$

where $\varrho$ is the density of the fluid, $\vec{u}$ the velocity vector, $\vec{n}$ the direction normal to the surface delimiting the control volume and $\overline{\overline{\tau}}$ the viscous stress tensor. We reorganize the right-hand side of eqn. 10 by considering the integrals on each boundary of the control volume separately (Fig. 6).

$$F_d = I_1 + I_2 + I_3 + I_4 \tag{11}$$

$$I_1 = -\rho \iint_1 (\vec{v} \cdot (-\overrightarrow{e_x}))\, u\, \mathrm{d}S \quad + \iint_1 p\, \mathrm{d}S \quad + \iint_1 -\tau_{xx}\, \mathrm{d}S \tag{12}$$

$$I_2 = -\rho \iint_2 (\vec{v} \cdot \overrightarrow{e_x})\, u\, \mathrm{d}S \quad - \iint_2 p\, \mathrm{d}S \quad + \iint_2 \tau_{xx}\, \mathrm{d}S \tag{13}$$

$$I_3 = -\rho \iint_3 (\vec{v} \cdot \overrightarrow{e_y})\, u\, \mathrm{d}S \quad + 0 \quad + \iint_3 \tau_{xy}\, \mathrm{d}S \tag{14}$$

$$I_4 = -\rho \iint_4 (\vec{v} \cdot (-\overrightarrow{e_y}))\, u\, \mathrm{d}S \quad + 0 \quad + \iint_4 -\tau_{xy}\, \mathrm{d}S \tag{15}$$

We then obtain

$$I_1 = +\rho \iint_1 u^2\, \mathrm{d}S \quad + \iint_1 p\, \mathrm{d}S \quad - 2\mu \iint_1 \frac{\partial u}{\partial x}\, \mathrm{d}S \tag{16}$$

$$I_2 = -\rho \iint_2 u^2\, \mathrm{d}S \quad - \iint_2 p\, \mathrm{d}S \quad + 2\mu \iint_2 \frac{\partial u}{\partial x}\, \mathrm{d}S \tag{17}$$

$$I_3 = -\rho \iint_3 uv\, \mathrm{d}S \quad + 0 \quad + \mu \iint_3 \frac{\partial u}{\partial y} + \frac{\partial v}{\partial x}\, \mathrm{d}S \tag{18}$$

$$I_4 = +\rho \iint_4 uv\, \mathrm{d}S \quad + 0 \quad - \mu \iint_4 \frac{\partial u}{\partial y} + \frac{\partial v}{\partial x}\, \mathrm{d}S \tag{19}$$

We extract the velocity components $u$ and $v$, the pressure $p$ and the derivatives $\frac{\partial u}{\partial x}$, $\frac{\partial u}{\partial y}$ and $\frac{\partial v}{\partial x}$ on each boundary of the control volume and process them with R (www.r-project.org) to calculate each term of the integrals. It should be noted that we assume that no parameter depends on the perpendicular coordinate $z$. As we also neglect border effects, integrating any variable such as drag per unit length in the $z$ direction is equivalent to multiplying the variable by the 10 cm-long length of the antenna in the $z$ direction.

Our PIV experiments are performed with a scaled model (scaled by a factor $\alpha$) in water (higher velocities) or oil (lower velocities). To obtain the corresponding drag in air, we used the drag coefficient. In case of equal Reynolds numbers, we have:

$$C_{d,\mathrm{air}} = C_{\mathrm{d,f}} \tag{20}$$

where the label $f$ refers to values in water/oil depending on the fluid used in the measurement. Eventually, we obtain:

$$F_{d,\mathrm{air}} = F_{d,\mathrm{f}} \frac{\rho_{\mathrm{air}}}{\rho_f} \frac{v^2_{\infty,\mathrm{air}}}{v^2_{\infty,f}} \frac{1}{\alpha^2} \tag{21}$$

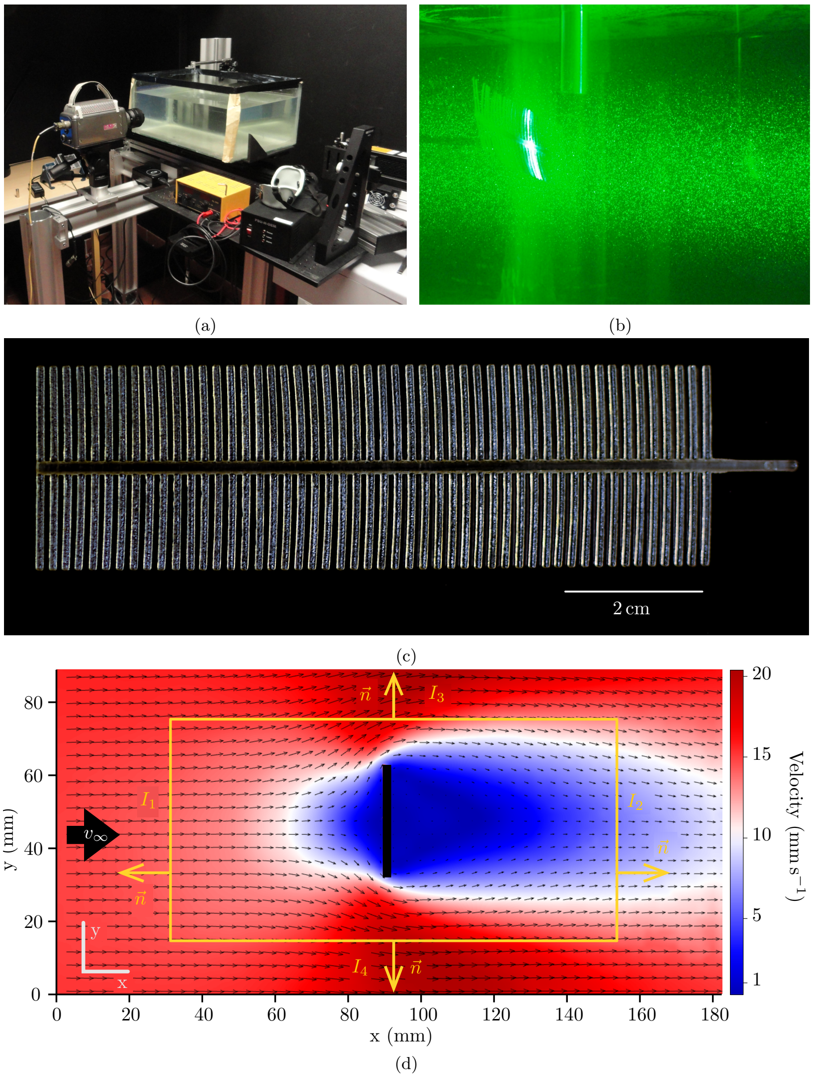


Figure 6: Experimental setup and definitions of the variables for the Particle Image Velocimetry (PIV) experiment. a: PIV setup. The tank is in the center and filled with water or oil depending on the simulated air velocity. The camera is on the left and the laser is positioned on the right. b: Laser sheet around the antenna enlightening the particles. c: 3D printed macrostructures. d: Control volume (orange) around an artificial antenna to extract the drag from PIV data. $\vec{n}$ is the vector normal to the control volume pointing outward. $I_1$, $I_2$, $I_3$ and $I_4$ are the integrals of the momentum through and the forces applied on each side of the control volume. $v_\infty$ is the far-field velocity. Here, the equivalent cylinders have a diameter of 105 µm. The artificial antenna is put in an oil tank with a velocity of 16 mm s$^{-1}$ corresponding to a far-field velocity of 0.05 m s$^{-1}$ in air.

#### Details of the Full Model WABBIT Simulations

In order to validate the method of pseudo-rami, we run computer-intensive simulations of the full antenna model (Fig. 7D), i.e. including flagellum, 100 rami and 28000 sensilla, using the open source Navier-Stokes solver 'WABBIT' (Wavelet-Adaptive Block-based solver for Insects in Turbulence) [6]. The code solves the incompressible Navier–Stokes equations using the artificial compressibility method, where a finite artificial speed of sound is introduced. We solve these governing equations using classical fourth order centered finite differences. The computational grid is dynamically adapted to the solution using Cohen-Daubechies-Feauveau wavelets (CDF4/0); this procedure refines the grid where it is necessary to achieve desired precision and coarsens it where possible, thus ensuring optimal exploitation of computational resources. Adaptation is done by locally refining the grid by a factor of two, where we allow for up to $J_{\max} = 11$ refinement steps. We use the volume penalization method to impose the no-slip condition at the fluid–solid interfaces. Parameters required for reproduction of numerical simulations are summarized in Tables 5 and 6.

A key property of turbulence is its multiscale nature. This property is similar, from a computational point of view, to the multiscale geometry of the insect pectinate antenna considered here, and our method is thus well suited for such simulations. Unlike in turbulent problems, viscosity plays a dominant role here, because the Reynolds number based on sensilla diameter and mean flow velocity is of the order of unity only. We therefore use the extensions to the WABBIT code developed for simulations of bristled wings [3, 7], most importantly the Runge-Kutta-Chebychev (RKC) schemes as described in the supplementary material to [3]. The RKC time integration schemes allow using larger time steps than the conventional Runge-Kutta schemes, and thus lowers the otherwise prohibitive cost of the numerical simulations. Detailed validation using numerical benchmarks [6, 3] as well as comparisons with experiments [4] demonstrate that the code produces reliable results.

In the present work, we use the 'fractal tree' module of the WABBIT code, which allows simulating the flow past an arbitrary arrangement of rigid, circular cylinders. Each cylinder segment is equipped with a semi-sphere at both ends to avoid sharp edges. The coordinates of cylinder start- and endpoints are computed using a custom python script prior to the actual simulation.

Even using our advanced numerical method, simulations of the complete antenna model remain challenging due to the vast difference of scales in the computation. We thus exploit the symmetry of the problem and simulate only one quarter of the complete problem, with an extension to WABBIT developed specifically for this article (Fig. 7A). Our computational domain is a cube of 4 cm with the antenna placed 1 cm downstream of the inlet (Fig. 7C). Free stream velocity is prescribed at the outer borders of the domain.

All simulations were run on the IRENE/ROME supercomputer hosted by the CEA in their TGCC computing center. Simulations used up to 4096 CPU cores and 8 TB of memory, and in total 1.966 million CPU hours were consumed for all simulations presented in this article. An individual simulation of the complete antenna model created a computational grid of about 300 000 blocks, each with $23^3$ individual grid points. The total number of grid points for a simulation was thus about 4 billion, yielding a total of 16 billion unknowns to be solved for in every time step. The simulations were run until the steady state was reached (Fig. 8).

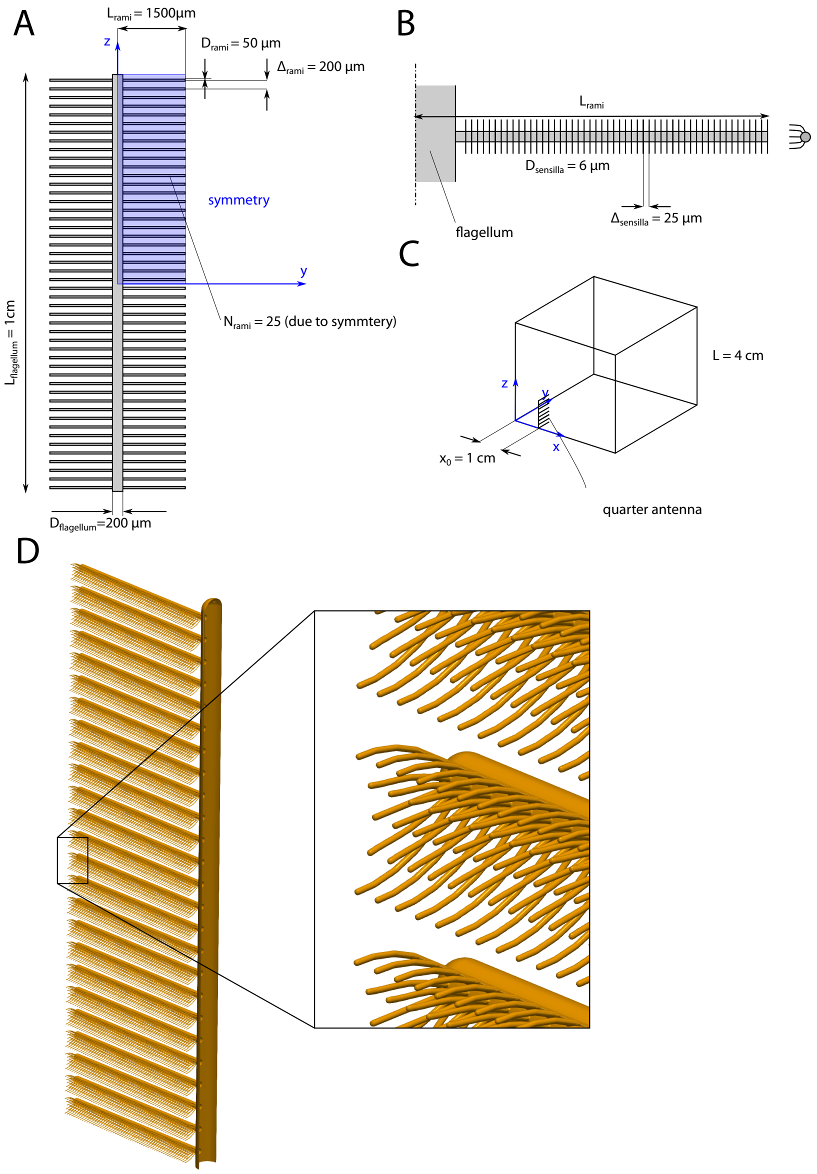


Figure 7: Key parameters for the numerical simulations of the full model of the pectinate antenna with the WABBIT code. (A) Sketch of the first two levels of geometrical structures (flagellum and rami). We simulate only the blue part of the antenna owing to symmetry. (B) Sketch of a single rami with sensillae. Inset shows a side view of the rami. (C) Sketch of the numerical setup with the coordinate system. Mean flow is in positive $x$-direction. (D) Rendering of the full model as it is used in the simulations.

| Parameter | Value | |
|---|---|---|
| | coarse | fine |
| Maximum number of refinements $J_{\max}$ | 10 | 11 |
| Grid points per rami diameter $(D_{rami}/\Delta x)$ | 28.2 | 56.3 |
| Grid points per sensilla diameter $(D_{sensilla}/\Delta x)$ | 3.4 | 6.8 |
| penalization parameter $K_\eta$ | 0.5 | |
| artificial speed of sound $C_0$ | 10 | |
| wavelet threshold $C_\varepsilon$ | $10^{-2}$ | |

Table 5: Full model simulations using WABBIT: numerical parameters.

| $u_\infty$ | resolution | CFL | RKC parameters [27, 3] |
|---|---|---|---|
| 1 m/s | coarse | 3.5 | $s = 10$, $\epsilon = 1.763$ |
| | fine | 3.45 | $s = 14$, $\epsilon = 1.495$ |
| 2 m/s | coarse | 2.84 | $s = 6$, $\epsilon = 0.958$ |
| | fine | 3.5 | $s = 10$, $\epsilon = 1.763$ |
| 5 m/s | coarse | 2.8 | $s = 4$, $\epsilon = 4.981$ |
| | fine | 1.0 | $s = 4$, $\epsilon = 20$ |

Table 6: Full model simulations using WABBIT: parameters for time integration. These parameters are critical for the cost of the numerical simulation and depend on the dimensionless viscosity (i.e. the mean flow velocity).

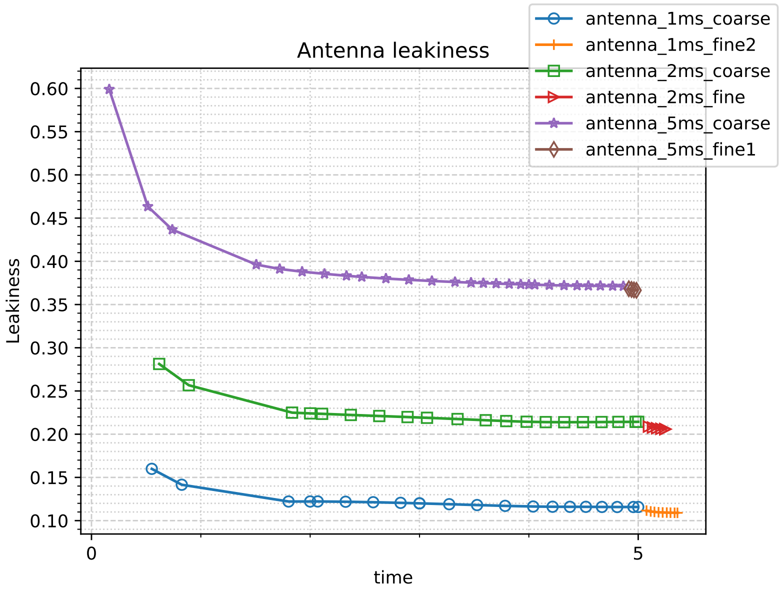


Figure 8: Temporal convergence of the simulations for the three velocities investigated here. The right part of the curves is calculated with a finer mesh than the left part.

#### Velocity field of the macrostructures

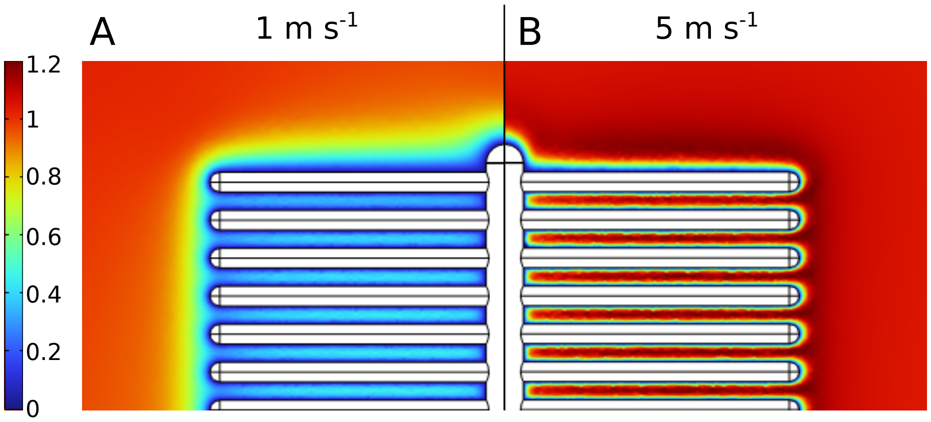


Figure 9: Normalized axial velocity ($u/u_\infty$) of the air passing through the macrostructure calculated with the simulations in Comsol. A: $u_\infty = 1\,\mathrm{m\,s^{-1}}$. B: $u_\infty = 5\,\mathrm{m\,s^{-1}}$.

#### Leakiness of a grid

To test the method of equivalent cylinders on another geometry, we considered a grid made out of two rows of cylinders perpendicular to each other. The inter-cylinder spacing is set at 200 µm and the cylinders all have a diameter of 50 µm. The grid is 1 cm by 3 mm. Using Comsol, we simulated a pattern (Fig 10, domain delimited by the red line) of the grid with periodic boundary conditions to simulate an infinite grid and calculated the equivalent diameter, equal to 46.3 µm.

We also simulated a quarter of the complete grid (Fig 11) as well as a row of 25 cylinders with a diameter of 46.3 µm, a length of 1.5 mm and an inter-cylinder spacing of 200 µm. In both cases, we used two symmetrical boundary conditions to simulate the entire structure. The far-field velocity was set at $1\,\mathrm{m\,s^{-1}}$.

Eventually, we found a leakiness of 18.7% for the grid, and 16.2% for the row of equivalent cylinders.

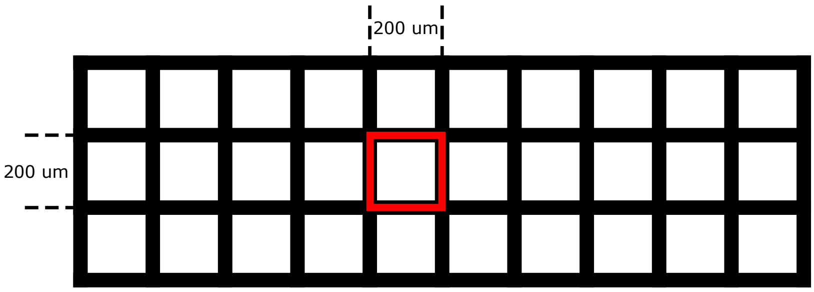


Figure 10: Geometry of the grid and pattern used to calculate the equivalent diameter.

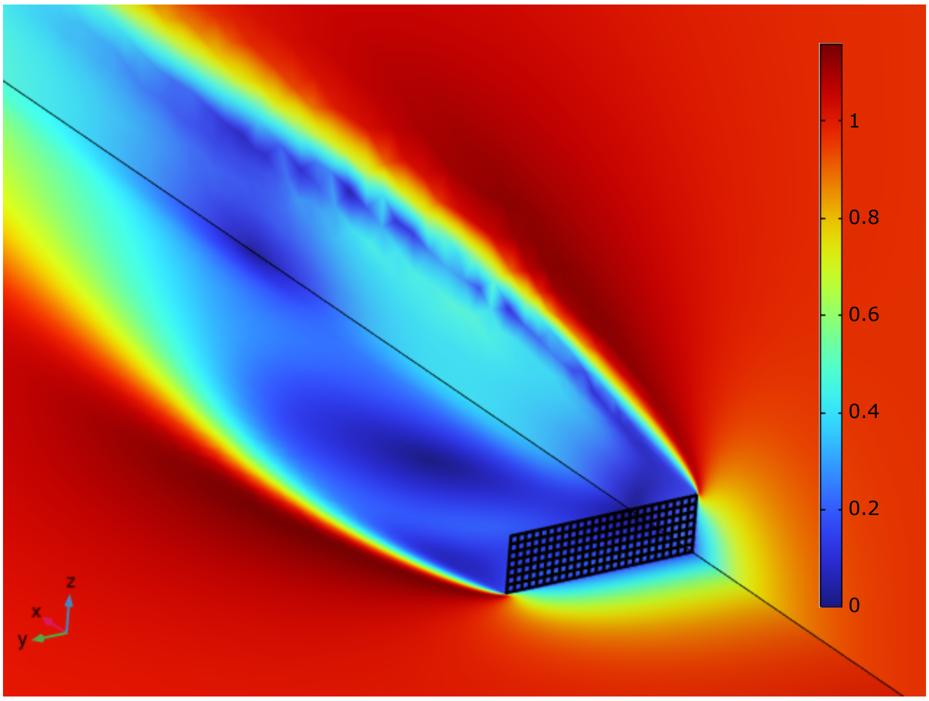


Figure 11: Simulation of the grid with Comsol. The far-field velocity is equal to $1\,\mathrm{m\,s^{-1}}$.

#### Comparison with the microstructure

To observe the influence of the rami and their sensilla with each other, we compare the leakiness of a single microstructure, measured in [5] with the one of the complete pectinate antenna calculated in this work. Leakiness of a complete antenna is lower than the one of the microstructure alone. It is thus useful to consider the entire antenna and not only the microstructure.

In our model, the distal half of a sensillum captures between 60% and 100% of the total amount of captured molecules (Fig. S12). This phenomenon of concentrated capture at the tip of the sensilla was already shown experimentally [15]. A passive model of mass transport on the microstructure (one ramus and its sensilla) [5] could explain this phenomenon. We show here that considering the entire antenna increases this gradient of capture between the proximal and basal parts of the sensilla and, hence, the amplitude of the olfactory lens phenomenon.

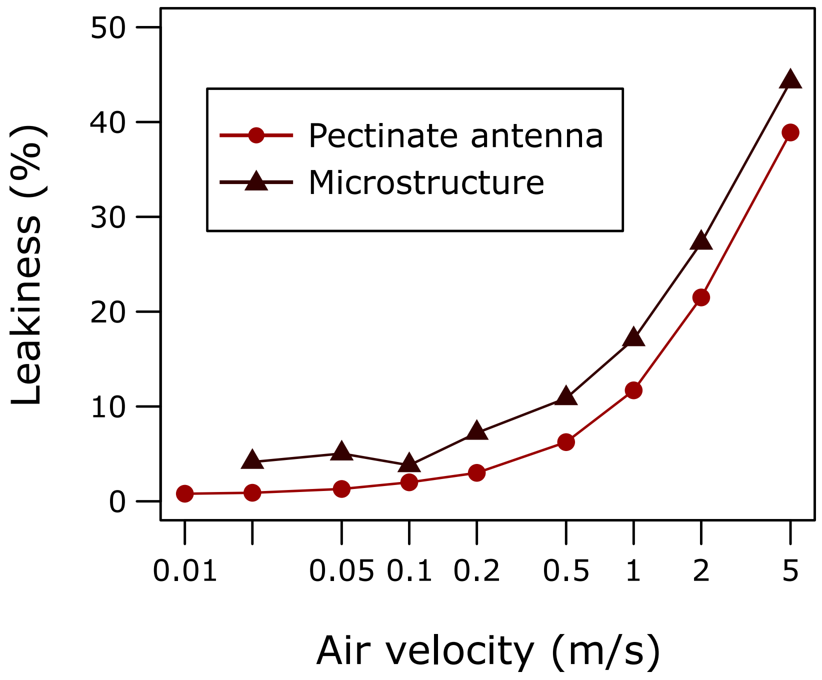


Figure 12: Leakiness of a complete pectinate antenna and of a microstructure alone [5] (black).

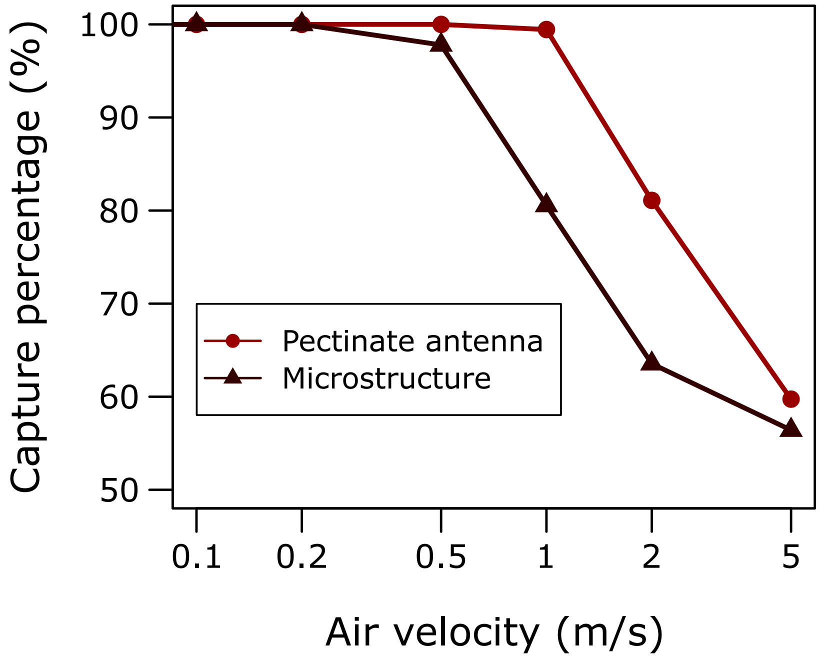


Figure 13: Relative capture by the distal half of the sensilla. The relative capture was calculated in the case of the study of the complete antenna (red) and compared with the case of the microstructure alone [5] (black).

### References


[1] T. Ando, S. Inomata, and M. Yamamoto. Lepidopteran Sex Pheromones. *Topics in Current Chemistry*, 239:51–96, 2004.

[2] Anonymous. 2020.

[3] Anonymous. 2020.

[4] Anonymous, 2020.

[5] Anonymous. 2020.

[6] Anonymous, 2021.

[7] Anonymous. 2021.

[8] R.S. Berger. Isolation, identification, and synthesis of the sex attractant of the cabbage looper, Trichoplusia ni. *Annals of the Entomological Society of America*, 59(4):767–771, 1966.

[9] A. Butenandt, R. Beckmann, and E. Hecker. Über den Sexuallockstoff des Seidenspinners, I. Der biologische Test und die Isolierung des reinen Sexuallockstoffes Bombykol. *Hoppe-Seyler s Zeitschrift für Physiologische Chemie*, 324(1):71–83, 1961.

[10] G. Gries, H. L. McBrien, R. Gries, J. Li, G. J. R. Judd, K. N. Slessor, J. H. Borden, R. F. Smith, M. Christie, J. T. Troubridge, P. D. C. Wimalaratne, and E. W. Underhill. (E4,E10)-dodecadienyl acetate: Novel sex pheromone component of tentiform leafminer, Phyllonorycter mespilella (Hübner) (Lepidoptera: Gracillariidae). *Journal of Chemical Ecology*, 19:1789–1798, 1993.

[11] R. Gries, D. Holden, G. Gries, P. D. C. Wimalaratne, K. N. Slessor, and C. Saunders. 3Z-cis-6,7-cis-9,10-Di-epoxy-heneicosene: Novel Class of Lepidopteran Pheromone. *Naturwissenschaften*, 84:219–221, may 1997.

[12] Regine Gries, Aurélia Reckziegel, Herman Bogenschütz, Hans-Günter Kontzog, Christian Schlegel, Wittko Francke, Jocelyn G. Millar, and Gerhard Gries. ( Z,Z )-11,13-Hexadecadienyl acetate and ( Z,E )-11,13,15-hexadecatrienyl acetate: synergistic sex pheromone components of oak processionary moth, Thaumetopoea processionea (Lepidoptera: Thaumetopoeidae). *Chemoecology*, 14:95–100, 2004.

[13] David R. Hall, Peter S. Beevor, Derek G. Campion, David J. Chamberlain, Alan Cork, Rosemary D. White, Aurelio Almestar, and Thomas J. Henneberry. Nitrate esters: Novel sex pheromone components of the cotton leafperforator, Bucculatrix thurberiella busck. (Lepidoptera: Lyonetiidae). *Tetrahedron Letters*, 33(33):4811–4814, aug 1992.

[14] H.E. Henderson, F.L. Warren, O.P.H. Augustyn, B.V. Burger, D.F. Schneider, P.R. Boshoff, H.S.C. Spies, and H. Geertsema. Isolation and structure of the sex-pheromone of the moth, Nudaurelia cytherea cytherea. *Journal of Insect Physiology*, 19(6):1257–1264, jun 1973.

[15] S. Kanaujia and K.-E. Kaissling. Interactions of pheromone with moth antennae: Adsorption, desorption and transport. *Journal of Insect Physiology*, 31(1):71–81, 1985.

[16] Michail V. Kozlov, Junwei Zhu, Peter Philipp, Wittko Francke, Elena L. Zvereva, Bill S. Hansson, and Christer Löfstedt. Pheromone specificity in Eriocrania semipurpurella (Stephens) and E. sangii (Wood) (Lepidoptera: Eriocraniidae) based on chirality of semiochemicals. *Journal of Chemical Ecology*, 22:431–454, mar 1996.

[17] D.F. Kurtulus, F. Scarano, and L. David. Unsteady aerodynamic forces estimation on a square cylinder by TR-PIV. *Experiments in Fluids*, 42(2):185–196, feb 2007.

[18] B. A. Leonhardt, V. C. Mastro, M. Schwarz, J. D. Tang, R. E. Charlton, A. Pellegrini-Toole, J. D. Warthen, C. P. Schwalbe, and R. T. Cardé. Identification of sex pheromone of browntail moth, Euproctis chrysorrhoea (L.) (Lepidoptera: Lymantriidae). *Journal of Chemical Ecology*, 17(5):897–910, may 1991.

[19] H. Ma and Z. Duan. Similarities of flow and heat transfer around a circular cylinder. *Symmetry*, 12(658):1–15, 2020.

[20] J. Millar, J. Steven McElfresh, and F.D.A. Marques. Unusual acetylenic sex pheromone of grape leaffolder (Lepidoptera: Pyralidae). *Journal of economic entomology*, 95(4):692–698, 2002.

[21] W. L. Roelofs, A. S. Hill, C. E. Linn, J. Meinwald, S. C. Jain, H. J. Herbert, and R. F. Smith. Sex Pheromone of the Winter Moth, a Geometrid with Unusually Low Temperature Precopulatory Responses. *Science*, 217(4560):657–659, aug 1982.

[22] Wendell L. Roelofs, Ada S. Hill, Ring T. Carde, and Thomas C. Baker. Two sex pheromone components of the tobacco budworm moth, Heliothis virescens. *Life Sciences*, 14(8):1555–1562, apr 1974.

[23] M.J. Tang, M. Shiraiwa, U. Pöschl, R.A. Cox, and M. Kalberer. Compilation and evaluation of gas phase diffusion coefficients of reactive trace gases in the atmosphere: Volume 2. Diffusivities of organic compounds, pressure-normalised mean free paths, and average Knudsen numbers for gas uptake calculations. *Atmospheric Chemistry and Physics*, 15(10):5585–5598, 2015.

[24] M. Tóth, H.R. Buser, A. Peña, H. Arn, K. Mori, T. Takeuchi, L.N. Nikolaeva, and B.G. Kovalev. Identification of (3Z,6Z)-1,3,6-9,10-epoxyheneicosatriene and (3Z,6Z)-1,3,6-9,10-epoxyeicosatriene in the sex pheromone of Hyphantria cunea. *Tetrahedron Letters*, 30(26):3405–3408, jan 1989.

[25] Miklós Tóth, Gábor Szöcs, Erik J. van Nieukerken, Peter Philipp, Frank Schmidt, and Wittko Francke. Novel type of sex pheromone structure identified from Stigmella malella (Stainton) (Lepidoptera: Nepticulidae). *Journal of Chemical Ecology*, 21:13–27, jan 1995.

[26] E. W. Underhill, J. G. Millar, R. A. Ring, J. W. Wong, D. Barton, and M. Giblin. Use of a sex attractant and an inhibitor for monitoring winter moth and bruce spanworm populations. *Journal of Chemical Ecology*, 13:1319–1330, jun 1987.

[27] J.G. Verwer, B.P. Sommeijer, and W. Hundsdorfer. RKC time-stepping for advection-diffusion-reaction problems. *Journal of Computational Physics*, 201(1):61–79, nov 2004.

[28] Sadao Wakamura, Norio Arakaki, Hiroshi Ono, and Hiroe Yasui. Identification of novel sex pheromone components from a tussock moth, Euproctis pulverea. *Entomologia Experimentalis et Applicata*, 100(1):109–117, jul 2001.

[29] John W. Wong, P. Palaniswamy, E. W. Underbill, W. F. Steck, and M. D. Chisholm. Novel sex pheromone components from the fall cankerworm moth, Alsophila pometaria. *Journal of Chemical Ecology*, 10:463–473, 1984.

[30] Hiroyuki Yamazawa, Naoto Nakajima, Sadao Wakamura, Norio Arakaki, Masanobu Yamamoto, and Tetsu Ando. Synthesis and Characterization of Diepoxyalkenes Derived from (3Z,6Z,9Z)-Trienes: Lymantriid Sex Pheromones and Their Candidates. *Journal of Chemical Ecology*, 27:2153–2167, 2001.

[31] Tetsuya Yasuda, Sachiko Yoshii, and Sadao Wakamura. Identification of the Sex Attractant Pheromone of the Browntail Moth, Euproctis similis (FUESSLY) (Lepidoptera: Lymantriidae). *Applied Entomology and Zoology*, 29(1):21–30, 1994.

[32] Junwei Zhu, Mikhail V. Kozlov, Peter Philipp, Wittko Francke, and Christer Löfstedt. Identification of a novel moth sex pheromone in Eriocrania cicatricella (Zett.) (Lepidoptera: Eriocraniidae) and its phylogenetic implications. *Journal of Chemical Ecology*, 21:29–43, jan 1995.
